# Discovery of protein modifications using high resolution differential mass spectrometry proteomics

**DOI:** 10.1101/2020.06.19.162321

**Authors:** Paolo Cifani, Zhi Li, Danmeng Luo, Mark Grivainis, Andrew M. Intlekofer, David Fenyö, Alex Kentsis

## Abstract

Recent studies have revealed diverse amino acid, post-translational and non-canonical modifications of proteins in diverse organisms and tissues. However, their unbiased detection and analysis remain hindered by technical limitations. Here, we present a spectral alignment method for the identification of protein modifications using high-resolution mass spectrometry proteomics. Termed SAMPEI for Spectral Alignment-based Modified PEptide Identification, this open-source algorithm is designed for the discovery of functional protein and peptide signaling modifications, without prior knowledge of their identities. Using synthetic standards and controlled chemical labeling experiments, we demonstrate its high specificity and sensitivity for the discovery of sub-stoichiometric protein modifications in complex cellular extracts. SAMPEI mapping of mouse macrophage differentiation revealed diverse post-translational protein modifications, including distinct forms of cysteine itaconatylation. SAMPEI’s robust parameterization and versatility are expected to facilitate the discovery of biological modifications of diverse macromolecules. SAMPEI is implemented as a Python package, and is available open-source from BioConda and GitHub (https://github.com/FenyoLab/SAMPEI).

## Introduction

Post-translational modifications (PTMs) control diverse biological processes, including activities of most cellular proteins. This involves reversible enzymatic modifications as part of biologic signaling, as well as various non-enzymatic chemical modifications during development, aging, and disease. As a result, identification of various protein modifications has propelled the discovery of numerous biological phenomena, molecular mechanisms, and disease processes.

Historically, PTM discovery followed the development of specific biochemical assays and affinity reagents, such as for example those for protein tyrosine phosphorylation (Aebersold et al. 2018). The development of high-resolution mass spectrometry provided a versatile and in principle unbiased approach for the identification of diverse protein modifications (Aebersold & Mann 2003; Aebersold & Mann 2016). This has revealed not only diverse protein PTMs, but also increasingly unanticipated modifications (Chick et al. 2015; Kong et al. 2017; Devabhaktuni et al. 2019). For example, recent mass spectrometry studies have reported lactylation and dopaminylation of histone residues in the regulation of gene expression (Zhang et al. 2019; Lepack et al. 2020). However, the extent and function of most protein modifications observed in biological tissues remain poorly defined, largely due to the technical challenges of their accurate and sensitive detection.

Most commonly, mass spectrometry proteomics is based on the analysis of proteolytic peptides (MacCoss et al. 2002; Mann & Jensen 2003; Smith et al. 2013). Tandem fragmentation of peptides and their corresponding high-resolution mass spectra, when sufficiently complete and unique, enable their identification and localization of distinct chemical modifications and substitutions. Most current approaches for peptide spectral matching (PSM) rely on statistical scoring of the similarity between the observed and theoretical spectra predicted based on their sequence and chemical modifications (Eng et al. 1994). In practice, this strategy limits the number of potential chemical modifications considered to a relatively small number of known adducts on specific residues, whose spectra are computed in modified and unmodified forms. Inclusion of multiple possible modifications expands the theoretical search space exponentially, thereby precluding the routine use of such approaches for unbiased PTM discovery. Indeed, in current experiments, most conventional methods for PSM assignment fail to match the majority of observed fragmentation spectra, at least in part due to diverse peptide modifications (Keller et al. 2002; Nielsen et al. 2006; Michalski et al. 2011).

Early attempts to circumvent this problem relied on the iterative analysis of unassigned spectra with different subsets of possible modifications (Jensen et al. 1997; Creasy & Cottrell 2002; Huang et al. 2013). However, only a relatively small number of known PTMs could be considered, with imprecise control of false discovery (Everett et al. 2010; Savitski et al. 2006). Restriction of the possible sequence search space using *de novo* sequence tags expanded the repertoire of variable PTMs that could be considered while significantly improving accuracy, but is restricted to spectra that have nearly complete fragment ion series (Mann & Wilm 1994; Searle et al. 2004; Na & Paek 2009; Dasari et al. 2010; Han et al. 2011). Search space reduction partially improved these limitations, but remains computationally expensive, as currently implemented in various algorithms (Bern et al. 2007; Bern & Kil 2011; Cox et al. 2011; Shortreed et al. 2015). Finally, recent implementations of fragment ion indexing and other efficiency concepts support fast assignment of modified mass spectra (Kong et al. 2017; Devabhaktuni et al. 2019; Bittremieux et al. 2019; Chi et al. 2018; Kahnert et al. 2020).

Sub-stoichiometric protein modifications that occur during biologic signaling, such as for example those induced by enzymes or metabolites, are particularly suited for identification using spectral alignment techniques, insofar as the sampling of both modified and unmodified states allows for their specific detection (Na et al. 2012; Pevzner et al. 2001; Tsur et al. 2005; Bandeira et al. 2007). Here, we extend this concept to develop SAMPEI, an open-source algorithm designed for the discovery of functional protein and peptide signaling modifications, without prior knowledge. We present its parameterization to optimize sensitivity and specificity using controlled chemical labeling experiments and synthetic peptides in complex cellular extracts, and demonstrate its utility by mapping protein signaling during mouse macrophage differentiation. Thus, SAMPEI and improved methods for mass spectrometry proteomics should enable comprehensive studies of protein signaling and chemical biology.

## Results

Differential display and spectral alignment have a long history in biology and physics for specific signal detection (Hokao & Francia 2001; Kentsis 2005). To extend this concept to regulatory protein signaling and chemical modifications, we reasoned that biologically functional protein modifications are present in modified and unmodified forms in different functional states. To implement this for mass spectrometric detection of differentially modified peptides, we developed the <u>s</u>pectral <u>a</u>lignment-based <u>m</u>odified <u>pep</u>tides <u>i</u>dentification (SAMPEI) algorithm. SAMPEI is based on the rationale that biologically regulated proteins with sub-stoichiometric modifications produce mass spectra from both the unmodified and modified peptides. These fragmentation spectra contain fragment ion series that are partly coincident and partly shifted by diagnostic mass shifts, corresponding to the masses of specific modifications (Zolg et al. 2018). To illustrate this concept, consider a model peptide DF_2_SAFINLVEF_11_CR with two possible modifications, either at the F_2_ or the F_11_ residues (Figure 1a). Collision-induced dissociation of F_2_-modified peptide is expected to produce fragment ion spectra with coincident y-ions, and a- and b-ions with specific mass shifts, corresponding to the F_2_ residue modification (Steen & Mann 2004). Conversely, modification of F_11_ residue is expected to produce y-ions with specific mass shifts as compared to those of the unmodified peptide, and the observable b- and a-ions that are largely coincident between modified and unmodified peptides (Figure 1A). Thus, given a set of high-resolution tandem mass spectra that have been matched to unmodified sequences, as obtained from current statistical database matching algorithms, SAMPEI performs pair-wise alignment of fragmentation mass spectra that are partly matched to unmodified sequences. This leads to the agnostic identification of possible PTMs based on the presence fragment ion series with specific alignment features (Figure 1b).

**Figure 1:**
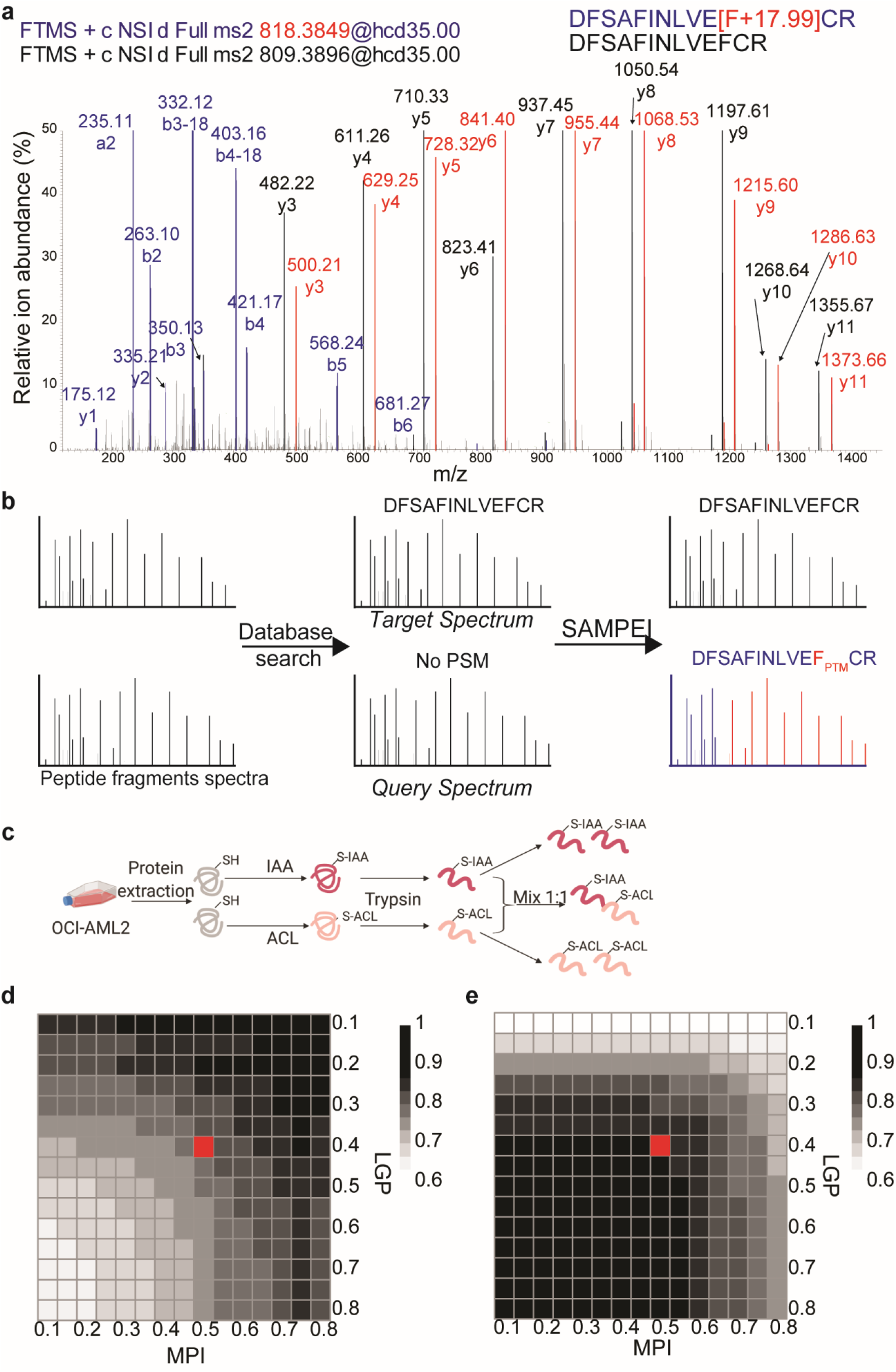
Outline and parameterization of Spectral Alignment-based Modified PEptide Identification (SAMPEI) algorithm, designed for agnostic discovery of peptide modifications. **a.** Overlay of specific alignment of fragment ions in high-resolution mass spectra of synthetic peptide DFSAFINLVEFCR unmodified (black) or containing fluorinated phenylalanine (blue and red, fluorination mass shift of +17.9905) at position 11. Fragment ions from the modified peptides that coincide with those from the unmodified form are marked in blue. Fragment ion mass shifts produced by phenylalanine fluorination are marked in red. **b.** Schematic of the SAMPEI algorithm: tandem mass spectra identified using peptide-spectral matching algorithms are used by SAMPEI as queries for partial matching against unmatched spectra, which often result from unpredicted chemical modifications and mutations. Based on the presence of partially coincident ion series, SAMPEI agnostically identifies sequences and mass shifts of modified peptides. **c.** Experimental schematic for the parameterization of sensitivity and specificity of SAMPEI using specific differentially alkylated proteomes. **d-e.** Heatmaps showing optimization of matched proportional intensity MPI and largest gap percentage LGP parameters to maximize precision (**d**) and recall (**e**), with carbamidomethylation as a known modification. Red symbols denote the default user-specified values (See also Figure S1).

We implemented SAMPEI using the Python programming language, openly available from Github using conda or pip (https://github.com/FenyoLab/SAMPEI). SAMPEI requires as input a set of high-resolution fragmentation spectra in MGF format and a list of high-confidence peptide-spectra matches (PSMs) defining a set of *target* spectra. By processing mass spectra in the open MGF format, SAMPEI is compatible with spectra recorded using all current high-resolution mass spectrometers and analyzed using most current algorithms for PSM assignment (Bern et al. 2007; Bern & Kil 2011; Cox et al. 2011; Shortreed et al. 2015). Tandem mass spectra are initially assigned to peptide sequences using any algorithm for peptide-spectral matching, and the resulting high-confidence PSMs define a set a of *query* spectra for SAMPEI. Here, we used X!Tandem for the initial implementation of SAMPEI, given its open source nature and ease of implementation on diverse computer systems, including high-performance cluster computers (Craig & Beavis 2004). Using X!Tandem expectation values, we selected high-confidence PSMs (1% false discovery rate) as the query spectra. High-confidence query PSMs were de. The remaining unmatched spectra were then considered by SAMPEI as candidates for agnostic identification of modified peptides, serving as *targets* for aligning the spectra of query PSMs. whose precursor ion falls within a user-defined mass tolerance window to all target fragment spectra. Users specify the molecular mass range of possible peptide modifications, which is set by default as less than 200 Da and greater than 10 Da. This range includes most currently known functional PTMs, as compiled in Unimod (http://www.unimod.org) (Creasy & Cottrell 2004), but excludes mass shifts from natural abundance isotopes. Each candidate target-query match is sequentially scored based on two orthogonal measures to evaluate their similarity. First, SAMPEI aligns discrete *m/z* ranges of the query and target spectra, and ranks putative matches based on the fraction of total fragment ion current measured in the query as compared to the target spectra, a metric defined as *matched proportional intensity* (MPI). Next, for each target-query match SAMPEI calculates the *largest gap percentage* (LGP) metric, defined as the fraction of the length of the peptide sequence consisting of consecutive undetected b- and y-ions. These parameters enable versatile control of accuracy and sensitivity, depending on the experimental goals.

We empirically calibrated these parameters to establish the sensitivity and specificity of SAMPEI for the agnostic discovery of specific peptide modifications in complex cellular proteomes in a controlled labeling experiment. We generated extracts of human OCI-AML2 cells and specifically modified their tryptic peptides with either iodoacetamide or acrylamide (Figure 1c). This generated otherwise identical complex mixtures with differentially alkylated peptides, bearing either 57.02 Da or 71.03 Da modifications, corresponding to cysteine carbamidomethylation or propionylation, respectively. Thus, we calculated the apparent sensitivity/recall and specificity/precision of detecting differentially alkylated peptides in complex mixtures as a function of variable MPI and LGP parameters. We used SAMPEI to compare high-resolution fragmentation spectra derived from human cell extracts containing peptides with either carbamidomethylated or propionylated cysteine residues, or their equimolar mixtures (Figure 1d-e; Supplementary Figure 1a-b). This allowed us to specify default MPI and LGP values that optimize SAMPEI sensitivity and specificity (marked by red symbols). These parameters can be adjusted by users to increase sensitivity or specificity, depending on the specific experimental needs. SAMPEI uses the fraction of matched target and query ion series to assign putative modifications to specific peptide residues. For spectra with incomplete ion series, putative modification site localization defaults to the first residue (N-terminus) in the peptide sequence. These features should allow for specific and sensitive detection of peptide modifications in complex cellular proteomes.

To test the ability of SAMPEI to discover unanticipated modifications, we designed a synthetic peptide bearing a specific chemical modification not observed naturally. From the analysis of currently described protein modifications curated in UniMod, we identified +17.99 Da as a unique mass shift not detected among any biological protein modifications thus far (Creasy & Cottrell 2004). Thus, we used the DF_2_SAFINLVEF_11_CR peptide derived from human PRKDC protein that is normally expressed by all human cells, and generated its two synthetic versions, containing fluorinated phenylalanine residues in position 2 or 11, expected to produce fragment ions series partly shifted by +17.99 Da due to fluorination. We confirmed the accurate composition of synthetic peptides by mass spectrometry in neat solvent and acquired high-resolution mass spectra of these peptides diluted in human OCI-AML2 cell extracts (Figure 2a; Supplementary Figure 2a). For both F_2_- and F_11_-fluorinated peptides, SAMPEI alignment correctly identified the +17.99 Da mass shift produced by this modification of the synthetic as compared to endogenous peptides. At the same time, spectra of fluorophenylalanine-containing peptides were not matched when the unmodified chemoform was absent in the queries pool, as modeled by a set of spectra recorded under identical experimental conditions from *E. coli* extracts (Figure 2b; Supplementary Figure 2b). These results indicate that SAMPEI is suitable for unbiased discovery of specific peptide modifications in complex biological proteomes.

**Fig. 2:**
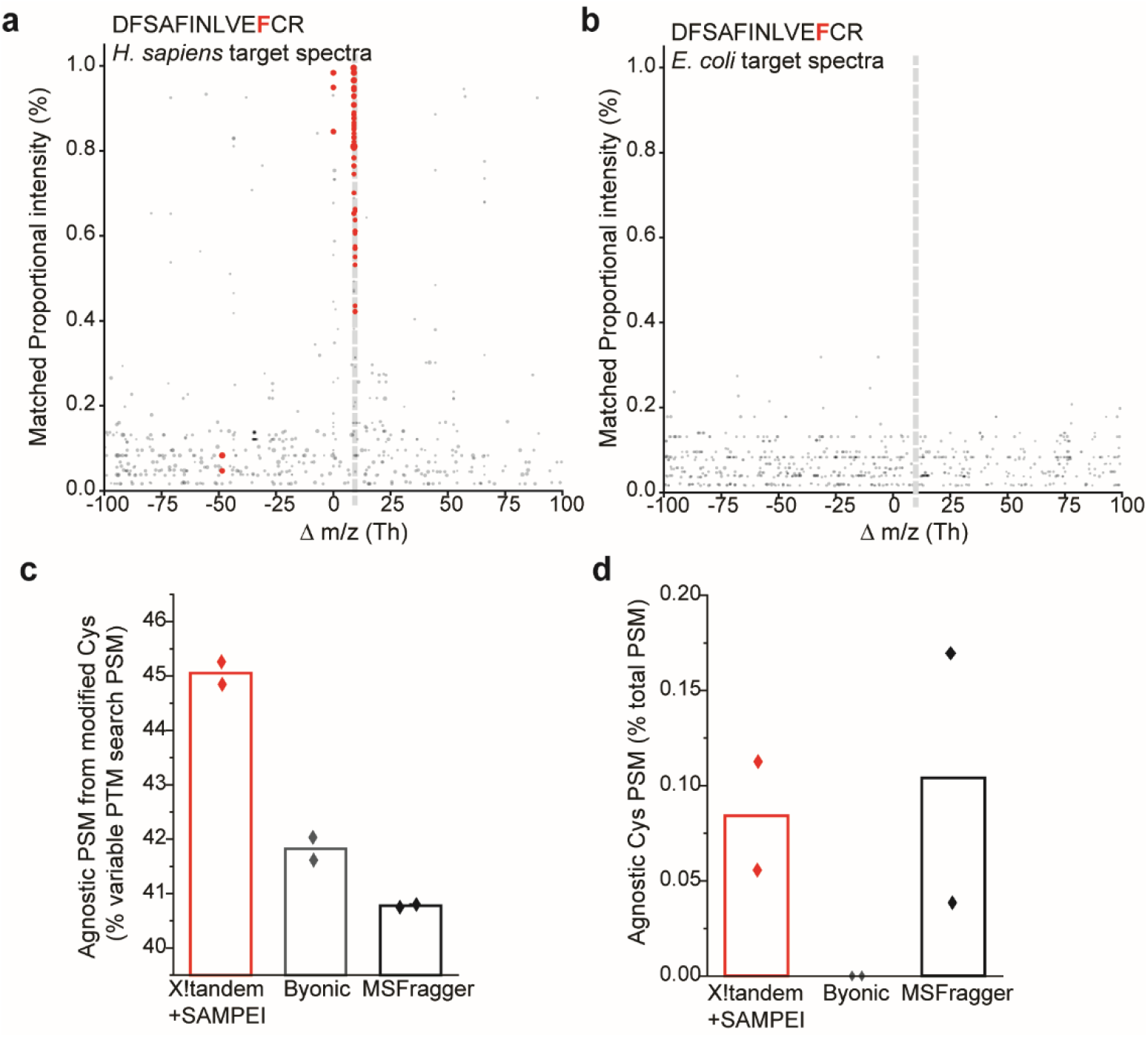
SAMPEI achieves sensitive and specific unbiased discovery of peptide modifications in complex cellular proteomes. **a,b.** Fraction of matched spectra as a function of modification mass shift (m/z) demonstrating accurate identification of synthetically fluorinated DFSAFINLVE(fluoro-F)CR peptide (17.99 Da mass shift, corresponding to 8.995 Th for +2 charged ions, as marked by dotted gray line) using as queries human cell extract containing the unmodified peptide (**a**) or an *E. coli* proteome not containing the unmodified chemoform, and thus serves as a negative control (**b**). Red symbols denote fluorinated peptide spectra matched by SAMPEI to the unmodified query PSM. Gray symbols denote matches of the fluorinated peptide with unrelated unmodified spectra (See also Figure S2). **c,d.** Comparison of sensitivity (**c**) and specificity (**d**) of agnostic identification of specific IAA or ACL cysteine modifications using SAMPEI, Byonic and MSFragger. Bars represent the mean values obtained for agnostic identification of carbamidomethylated and propionylated cysteines respectively. For each experiment, the two symbols represent the values obtained in two experiments, where cysteine fixed modification was set either to carbamidomethylation or propionylation (See also Figure S3).

To establish the relative sensitivity and specificity of SAMPEI, we compared its performance with MSFragger and Byonic algorithms that are compatible with open format spectral analysis for unbiased PTM identification (Kong et al. 2017; Bern et al. 2007; Bern & Kil 2011). We analyzed human OCI-AML2 cell extracts containing differentially carbamidomethylated or propionylated cysteine residues, converted the RAW files into MGF format using uniform parameters, and analyzed them using SAMPEI, MSFragger, and Byonic to agnostically detect differentially specific cysteine alkylation (the experimental design is shown in Figure 1c). To determine the relative sensitivity and specificity, we analyzed the spectra setting either cysteine carbamidomethylation or propionylation as a fixed modification and calculating the agnostic recovery of the alternative modification. We observed that the fraction of PSMs from specifically modified peptides detected by SAMPEI was slightly greater than those calculated by both MSFragger and Byonic, which is a measure of sensitivity (45% versus 41% and 42%, respectively; Figure 2c, Supplementary Figure 3). We reasoned that spectra produced by peptides with cysteines that are exclusively carbamidomethylated or propionylated can be used to assess the specificity of agnostic detection of peptide modifications. Thus, we defined PSMs with apparent cysteine propionylation in samples labeled solely with iodoacetamide (where carbamidomethylation was expected) as false identifications, and vice versa. This analysis demonstrated that the default parameterization of SAMPEI has a mean false unbiased identification rate for modified peptides in complex human cell extracts of 0.09%, as compared to 0.1% and 0% by MSFragger and Byonic, respectively (Figure 2d). Thus, SAMPEI demonstrates excellent specificity and sensitivity for the unbiased discovery of protein modifications in complex proteomes.

To establish the relative sensitivity and specificity of SAMPEI, we compared its performance with MSFragger and Byonic algorithms that are compatible with open format spectral analysis for unbiased PTM identification (Kong et al. 2017; Bern et al. 2007; Bern & Kil 2011). We analyzed human OCI-AML2 cell extracts containing differentially carbamidomethylated or propionylated cysteine residues, converted the RAW files into MGF format using uniform parameters, and analyzed them using SAMPEI, MSFragger, and Byonic to agnostically detect differentially specific cysteine alkylation (the experimental design is shown in Figure 1c). To determine the relative sensitivity and specificity, we analyzed the spectra setting either cysteine carbamidomethylation or propionylation as a fixed modification and calculating the agnostic recovery of the alternative modification. We observed that the fraction of PSMs from specifically modified peptides detected by SAMPEI was slightly greater than those calculated by both MSFragger and Byonic, which is a measure of sensitivity (45% versus 41% and 42%, respectively; Figure 2c, Supplementary Figure 3). We reasoned that spectra produced by peptides with cysteines that are exclusively carbamidomethylated or propionylated can be used to assess the specificity of agnostic detection of peptide modifications. Thus, we defined PSMs with apparent cysteine propionylation in samples labeled solely with iodoacetamide (where carbamidomethylation was expected) as false identifications, and vice versa. This analysis demonstrated that the default parameterization of SAMPEI has a mean false unbiased identification rate for modified peptides in complex human cell extracts of 0.09%, as compared to 0.1% and 0% by MSFragger and Byonic, respectively (Figure 2d). Thus, SAMPEI demonstrates excellent specificity and sensitivity for the unbiased discovery of protein modifications in complex proteomes.

SAMPEI is motivated by the hypothesis that unanticipated regulatory protein modifications can be discovered from comprehensive analysis of biological systems. To explore this idea, we sought to map protein modifications induced during mammalian cell differentiation. We chose to study the response of mouse RAW264.7 macrophage cells to lipopolysaccharide (LPS), a potent inducer of macrophage activation and differentiation that involves extensive protein and metabolic signaling (Alasoo et al. 2015; Kamal et al. 2018; Rattigan et al. 2018; Seim et al. 2019). We treated RAW264.7 cells with LPS using established methods and analyzed the resultant proteomes by two-dimensional nanoscale liquid chromatography and high-resolution mass spectrometry proteomics (Cifani et al. 2018). Using SAMPEI parameters designed to maximize the specificity of unbiased discovery (*MPI* ≥ 0.5, *LGP* ≤ 0.4), we identified 21,846 unique PSMs with putative peptide modifications producing mass shift alignments between 10 and 200 Da (Figure 3b, Supplementary Figure 4). SAMPEI assigned approximately 60% of putative modifications to specific peptide residues, with the remainder assigned to the N-terminal residue by default. Putative modifications identified by SAMPEI include several commonly observed ones, such as methionine oxidation (+16 Da), lysine acetylation (+42 Da) and serine and threonine phosphorylation (+80 Da). Consistent with the known reactivity of high concentrations of iodoacetamide as part of sample preparation, we also observed putative carbamidomethylation (+57 Da) of non-cysteine residues, in agreement with prior reports (Chick et al. 2015; Devabhaktuni et al. 2019). Importantly, SAMPEI also identified peptide modifications with a wide range of diverse mass shifts (Figure 3c, Supplementary Figure 4). It is probable that many apparent amino acid modifications, in particular those on aliphatic residues, represent amino acid substitutions due to somatic nucleotide mutations acquired in cultured cells, as recently documented using proteogenomic approaches (Cifani et al. 2018). Many of the putative modifications have also been observed in prior studies, including transpeptidation, and chemical transformations of various amino acid side chains by glycans and ion coordination (Kong et al. 2017; Devabhaktuni et al. 2019). While some of them arise from biochemical reactions upon cell lysis and proteome processing, we anticipate that many of them occur biologically and represent unanticipated regulatory processes.

**Fig. 3:**
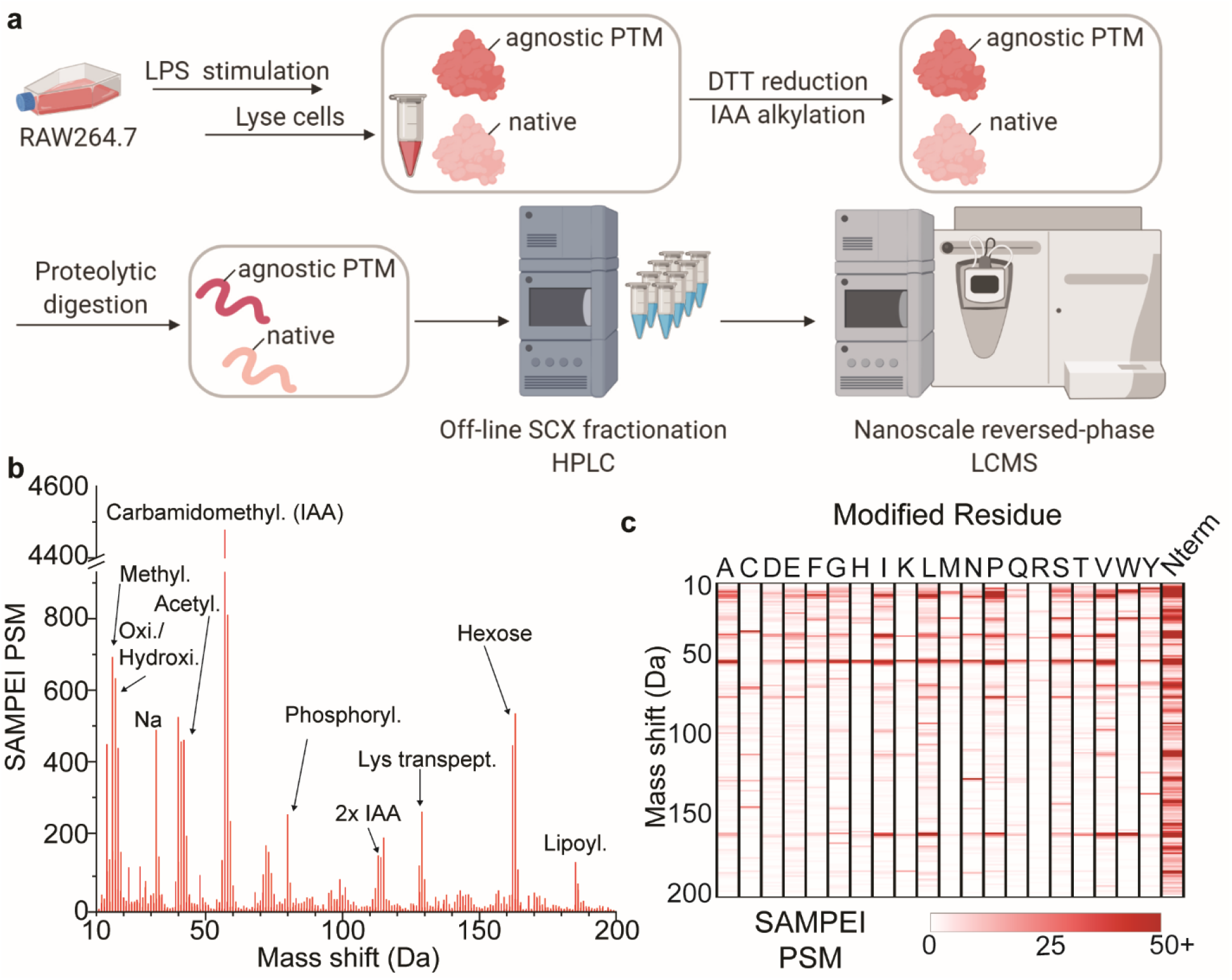
Agnostic protein signaling discovery in LPS-stimulated macrophage differentiation. **a.** Schematic of RAW264.7 macrophage cells LPS-induced differentiation, followed by proteome extraction, proteolysis and peptide chromatography and high-resolution tandem mass spectrometry. **b.** Histogram of peptide-spectrum matches (PSM) as a function of peptide modification mass shifts (21,846 peptides with MPI ≥ 0.7). Selected spectral matches with putative modifications are labeled with arrows. **c.** Frequency and putative amino acid localization of chemical adduct molecular weights discovered by SAMPEI (See also Figure S4).

The known reactivity of cysteine residues in proteins led us to examine their modifications upon LPS-induced macrophage activation (Poole & Nelson 2008; Seo & Carroll 2009; Jeong et al. 2011). Thus, we examined the most abundant putative modifications assigned by SAMPEI to cysteine residues upon LPS treatment (Figure 4a). We attempted to deduce their identity from the analysis of high-accuracy mass measurements, as additionally modified by single or double carbamidomethylation induced by iodoacetamide. While some of the observed mass shift alignments could be attributed to known modifications annotated by UniMod, we also observed unanticipated +130.03 Da and +146.02 Da mass shifts (Figure 4a). We confirmed these SAMPEI identifications by manual inspection of the corresponding high-resolution fragment ion spectra from several modified and unmodified peptide pairs (Figure 4b-c, Supplementary Figures 5–6). The observed high-accuracy mass measurements of the putative modifications are consistent with the C_5_H_6_O_4_ and C_5_H_6_O_5_ elemental composition for the +130.03 Da and +146.02 Da modifications, respectively.

**Fig. 4:**
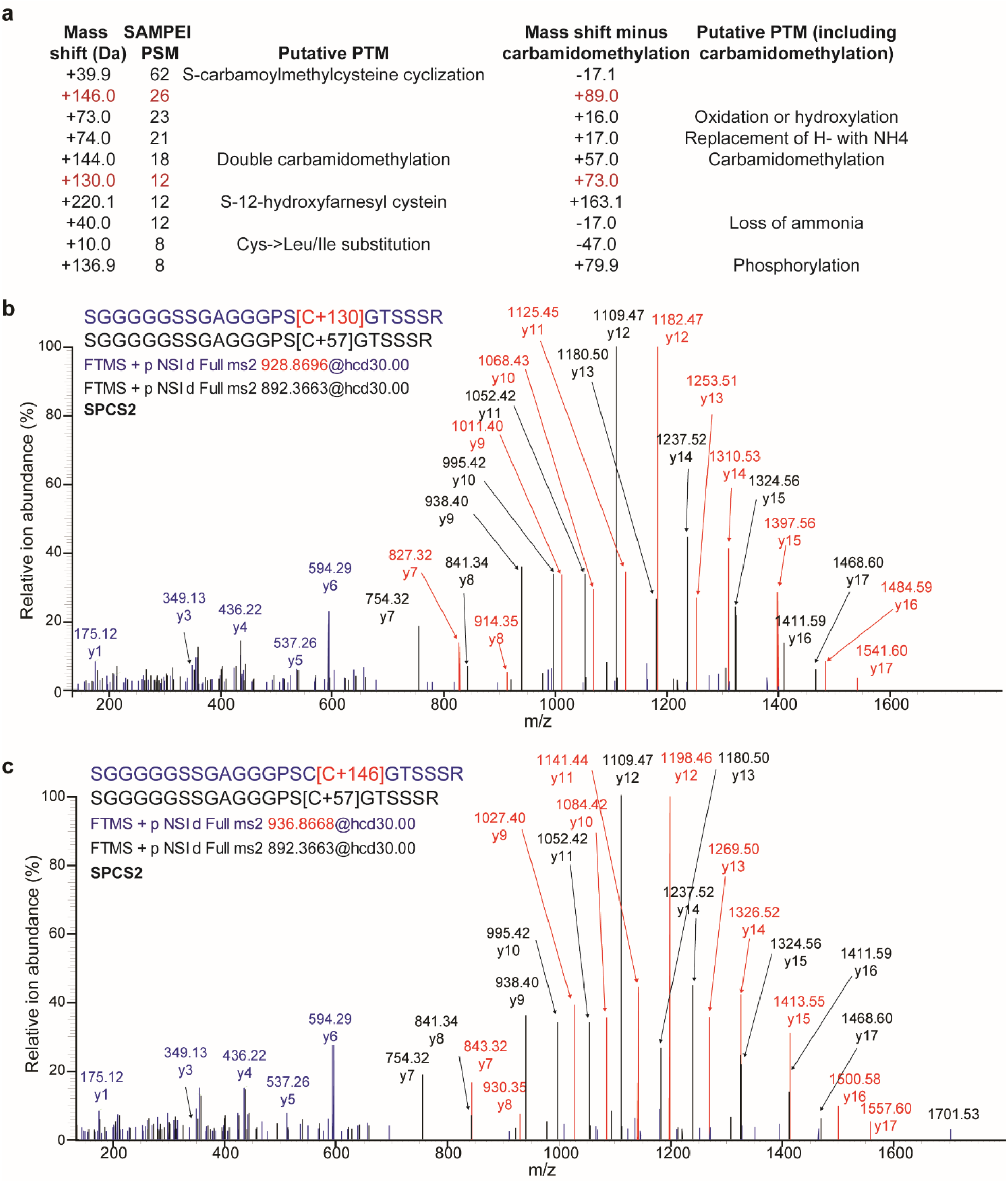
Non-canonical cysteine modifications discovered by SAMPEI. **a.** Ten most prevalent non-canonical cysteine PTMs discovered by SAMPEI, with annotation of putative adducts based on high-accuracy mass measurements. Concurrent carbamidomethylation was considered in the two right-most columns. Red denotes modification molecular weights not annotated in UniMod. **b-c.** Representative mass spectra pairs of Spcs2 peptides from containing either unmodified, experimentally carbamidomethylated cysteine, or bearing 130.03 Da (b) and 146.02 Da (c) modifications. Ions shared by unmodified and modified peptides (blue), those exclusive for unmodified peptides (black), and those specific for +130.03 Da or +146.02 Da modified peptides (red) are labelled accordingly (See also Figure S5–S6).

Recently, LPS-stimulated macrophages were reported to produce itaconate, an electrophilic dicarboxylic acid that is similar to succinate and fumarate TCA cycle intermediates. Itaconate can chemically react with nucleophilic amino acids (Strelko et al. 2011; Cordes et al. 2016; Kulkarni et al. 2019; Mills et al. 2018). Indeed, we confirmed the generation of free itaconate and the expression of Acod1 (Irg1) aconitate decarboxylase enzyme responsible for its production in RAW264.7 cells upon LPS stimulation using small molecule mass spectrometry and Western immunoblotting, respectively (Figure 5a-c, Supplementary Figure 7). Gene Ontology (GO) annotation of all unique peptides with putative itaconate modifications revealed enrichment for mitochondrion and ribosome localization, and for processes related to glycolysis, protein translation and stabilization (Supplementary Figure 8). Interestingly, we observed cysteine itaconatylation of peptides corresponding to gamma-interferon-inducible lysosomal thiol reductase (Gilt) and protein disulfide-isomerase A3 (Pdia3), both of which regulate cellular thiol/disulfide redox balance. We also observed itaconatylation of Gapdh and Aldoa, similar to a recent study of thiol reactivity using chemical profiling (Qin et al. 2019). While the +130.03 Da modified proteins can be explained by itaconate, we reasoned that the +146.02 Da modified peptides may be due to the modification by oxidized itaconate or oxidation of itaconatylated cysteine to sulfoxide, an oxidized +15.99 Da form of cysteine induced by redox cellular signaling. Given that redox signaling can also contribute to macrophage activation (West et al. 2011; Paulsen & Carroll 2013), these results suggest that LPS-induced mouse macrophage differentiation involves cysteine-dependent signaling by apparent itaconatylation and oxidation of cysteine.

**Fig. 5:**
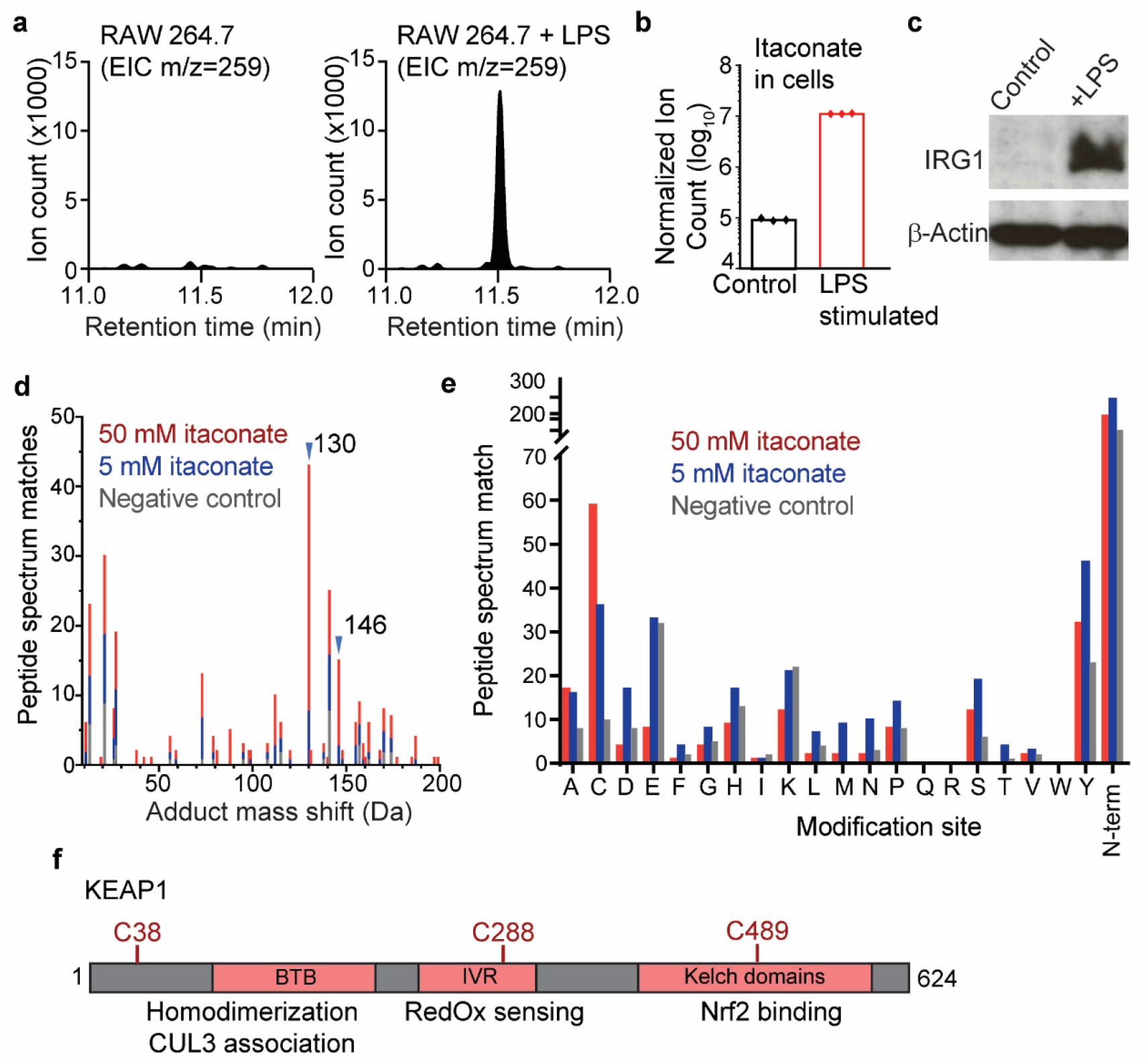
Validation of itaconate-induced cysteine modifications. **a.** RAW 264.7 macrophages were unstimulated (left) or stimulated with LPS (right), followed by metabolite extraction and GC-MS analysis. Extracted ion chromatograms (EIC) for m/z 259 are shown, demonstrating induction of itaconate in response to LPS stimulation (See also Figure S9). **b.** Quantification of itaconate production by RAW264.7 macrophages upon LPS stimulation as measured by LC-MS in cells (*n* = 3 biologic replicates). **c**. Representative Western blot of Irg1 protein expression in RAW264.7 macrophages upon LPS-stimulation, with actin as loading control. **d.** Itaconate reacts with purified BSA and KEAP1 *in vitro* to produce specific cysteine modifications, as shown by the distribution of identified spectra as a function of increasing itaconate concentration (pink and red), as compared to PBS control (gray). Blue arrowheads mark concentration-dependent induction of +130 Da and +146 Da modified peptides (See also Figure S10,S11,S12). **e.** Putative site localization of observed modifications in BSA and KEAP1, with specific itaconate induction of cysteine modifications. **f.** Schematic representation of KEAP1 protein functional domains, with itaconate modified cysteine residues.

To test this hypothesis and directly confirm protein cysteine itaconatylation detected by SAMPEI, we examined purified recombinant proteins upon exposure to physiologically relevant concentrations of itaconate in vitro. We chose BSA given its established suitability for mass spectrometry analysis, as well as KEAP1, a known protein sensor of cellular redox homeostasis (Sihvola & Levonen 2017). We treated purified BSA and KEAP1 proteins with either 5 or 50 mM synthetic itaconate, as compared to mock control reactions, and analyzed the resultant chemical products by independent analysis using digestions with trypsin, GluC, and elastase proteases, followed by high-resolution tandem mass spectrometry (Supplementary Figure 9). As predicted, we observed concentration-dependent formation of both +130.03 Da and +146.02 Da modifications on cysteine residues, specifically in the presence of itaconate but not in mock control treated reactions (Figure 5d-e, Supplementary Figure 10-12). We confirmed that the two unanticipated cysteine modifications identified in LPS-activated macrophage were indeed due to itaconate. Specifically, we detected four KEAP1 peptides modified with the +130.0 Da adducts on cysteine residues 38, 288, and 489 (Figure 5f). While alkylation of Cys288 was previously observed (Mills et al. 2018), Cys38 and Cys489 represent additional regulatory alkylation sites. We verified the presence of identical itaconatylated cysteine residues in independent, differentially digested experimental replicates of both BSA and KEAP1, such as for example Cys489 itaconatylation in both trypsin and elastase-digested samples. Thus, physiologically relevant concentrations of itaconate can induce cysteine itaconatylation in diverse proteins.

## Discussion

Covalent modifications of diverse macromolecules contribute to multiple forms of biological signaling. In the case of protein signaling, recent studies have revealed an increasingly diverse and unanticipated repertoire of enzymatic and non-enzymatic amino acid modifications. The ability to comprehensively map and identify protein modifications is critical to understand biochemical mechanisms of biological signaling. SAMPEI is designed for the identification of peptide modifications from high-resolution mass spectrometry data without prior knowledge.

The advantage of SAMPEI is its explicit detection of pairs of modified and unmodified peptides, thereby enabling the identification of potential biochemical signaling events that occur differentially in distinct biological states. Globally, SAMPEI exhibits excellent sensitivity that is on par with open database searches, while maintaining high specificity. We anticipate that extension of future versions of SAMPEI to incorporate high-resolution MS3 fragmentation spectra will enable direct analysis of the composition amino acid adducts and non-canonical amino acids, as enabled by the development of small molecule spectral libraries such as METLIN (Guijas et al. 2018). While we used X!Tandem for the identification of unmodified peptides, SAMPEI is compatible with all currently used peptide spectral matching algorithms, including open format input.

We expect that improved peptide fragmentation methods can be used to boost the sensitivity of protein modification mapping by SAMPEI by generating informative ion series that contain chemical modifications that currently alter the properties of collision-induced peptide fragmentation. Similarly, improved scoring functions can be integrated to enhance the detection of multiple combined modifications. In addition, the use of proteogenomic strategies for the initial database-driven identification of unmodified peptides can be used to assign spectra with mass shifts produced by amino acid substitutions (Cifani et al. 2018), thereby improving the sensitivity and specificity of PTM identifications. SAMPEI can be directly integrated with programs for quantitative analysis such as MaxQuant (Cox & Mann 2008), which can be used for precise studies of biological PTM dynamics. For example, quantitation of putative PTMs can aid in the identification of mechanisms of their regulation (Kahnert et al. 2020).

Most importantly, SAMPEI and other methods designed for PTM identification without prior knowledge are ideally suited for biological discovery. We demonstrate this by the identification of unanticipated forms of protein cysteine itaconatylation during LPS-induced macrophage activation, as confirmed by direct labeling experiments. Some of the itaconatylated proteins induced upon macrophage activation are involved in cellular redox homeostasis, suggesting a mechanistic link between TCA cycle reprogramming and redox signaling, as supported by the identification of oxidized forms of itaconatylated cysteine upon macrophage activation. Further studies will be needed to define the mechanisms and functions of this unanticipated signaling interaction. Similarly, future studies will be needed to define the contribution of KEAP1 Cys38 and Cys489 oxidation and itaconatylation to its control of macrophage function.

The extent and function of most protein modifications observed in biological tissues remain largely undefined, largely due to the technical challenges of their accurate and sensitive detection. In addition to enzymatically derived PTMs, recent studies have also revealed protein modifications produced by the spontaneous reaction of cellular metabolites (Zheng et al. 2020; Qin et al. 2020). Enzymatic and metabolite-derived PTMs may signal by regulating enzymatic activities or protein interactions, as exemplified by the direct link between energetic metabolites and post-translational histone modifications in cells (Gut & Verdin 2013). As sensitivity of mass spectrometry now approaches near-complete direct analysis of complex cellular and tissue proteomes (Wilhelm et al. 2014; Zhou et al. 2013), improved methods for the discovery of protein and macromolecular modifications without prior knowledge, such as SAMPEI, should enable comprehensive studies of biological signaling.

## Acknowledgements

We thank Henrik Molina, Soeren Heissel, and Caitlin Streckler for technical assistance. This research was supported by the NIH R01 CA204396, R01 CA214812, R21 CA235285, P30 CA008748, St. Baldrick’s Foundation, Hyundai Hope on Wheels, Burroughs Wellcome Fund, Damon Runyon-Richard Lumsden Foundation, Rita Allen Foundation, Leukemia & Lymphoma Society, the Starr Cancer Consortium, Pershing Square Sohn and Mathers Foundations, and Mr. William H. and Mrs. Alice Goodwin and the Commonwealth Foundation for Cancer Research and the Center for Experimental Therapeutics at MSKCC.

## Author Contributions

Conceptualization, AK, PC, DL, DF; Investigation, PC, DL, ZL, MG, AI; Analysis, DL, PC, ZL, DF, AK; Resources, DF, AK; Writing of original draft, PC, DL, AK; Writing of final draft, all authors; Funding acquisition, DF, AK.

## STAR Methods

### Reagents

Iodoacetamide (IAA, ≥99%, NMR grade), acrylamide (ACL, certified reference grade), itaconate (ITA, ≥96.0%, analytical grade) and lipopolysaccharides (LPS, from Escherichia coli O111:B4), guanidinium chloride, ammonium bicarbonate (ABC), dithiothreitol (DTT) were obtained from Sigma Aldrich. LC-MS grade H_2_O and acetonitrile (ACN) were from Fisher Scientific. LC-MS grade formic acid (FA) and trifluoroacetic acid (TFA) of >99% purity was obtained from Thermo Scientific. LysC (mass spectrometry grade) was obtained from FUJIFILM Wako Chemicals. Trypsin (modified, sequencing grade), GluC (sequencing grade) and elastase were obtained from Promega. Bovine serum albumin (BSA) was obtained from Sigma Aldrich. Human recombinant Kelch-like ECH-associated protein 1 (KEAP1, His & GST Tag) was obtained from Sino Biological, and the lyophilized formulation was prepared in 20 mM Tris, 500 mM NaCl, 10% glycerol, pH 7.4, 5% trehalose, 5% mannitol. Protease inhibitors 4-(2-aminoethyl)-benzenesulfonyl fluoride hydrochloride (AEBSF) and pepstatin were obtained from Santa Cruz, bestatin from Alfa Aesar, and leupeptin from EMD Millipore.

Synthetic DFSAFILVEFCR peptides were produced by solid phase synthesis and purified to 95% purity by liquid chromatography (New England Peptides) in three chemoforms, all with carbamidomethylated C_12_: 1) with unmodified phenylalanines, 2) with fluoro-F_2_, and 3) with fluoro-F_11_. Peptide composition and chemical modifications were confirmed using mass spectrometry.

### Cell culture

Human OCI-AML2 and mouse RAW264.7 cells were obtained from the American Type Culture Collection (ATCC). The identity of cells was verified by STR analysis (MSKCC Integrated Genomics Operation). The absence of Mycoplasma sp. contamination was determined using Lonza MycoAlert (Lonza). OCI-AML2 cells were cultured in RPMI 1640 (Corning) supplemented with 10% fetal bovine serum (FBS), 1% penicillin/streptomycin and 1% L-glutamine. RAW264.7cellls were cultured in DMEM with sodium pyruvate (Corning) supplemented with 10% fetal bovine serum (FBS) and 1% penicillin/streptomycin. Cells were cultured in 5% CO_2_ in a humidified atmosphere at 37 °C.

### Preparation of OCI-AML2 proteome alkylated with iodoacetamide (IAA) or acrylamide (ACL)

4×10^6^ of OCI-AML2 cells were pelleted by centrifugation at 400 G for 5 min at 4 °C. The cell pellets were washed twice with PBS, resuspended in cold lysis buffer (6 M guanidinium chloride, 100 mM ABC, pH 8.4), and lysed using S220 adaptive focused sonicator (Covaris). Protein concentration was determined using the bicinchoninic acid (BCA) assay according to the manufacturer’s instructions (Pierce). Proteomes were reduced by dithiothreitol (DTT, 10 mM) at 56°C for 45 min and then divided into two equal fractions, and alkylated by 55 mM iodoacetamide (IAA, 55 mM) and acrylamide (ACL, 55 mM), respectively (room temperature, 30 min in the dark). Alkylation was quenched by the addition of DTT (10 mM, 56°C, 30 min). Lysates were diluted with 100 mM aqueous ABC, pH 8.4, to a final concentration of 0.6 M guanidinium chloride and the proteomes were proteolyzed using 1:100 w/w (protease:proteome) LysC endopeptidase at 37°C for 6 h. The solution was further diluted with 100 mM aqueous ABC to 0.3 M guanidinium chloride, and the proteomes were digested using 1:50 w/w modified trypsin at 37°C overnight. Digestion was stopped by acidifying the reactions to pH 3 using aqueous trifluoroacetic acid (TFA), peptides were desalted using solid-phase extraction with C18 MacroSpin columns (Nest Group), and lyophilized by vacuum centrifugation (Genevac EZ-2 Elite).

### Preparation of RAW264.7 proteome under LPS stimulation

RAW264.7 cells were seeded 10×10^6^ cells per 150 mm dish (6 dishes), and allowed to adhere for 24 h. LPS was added to the medium at 50 ng/mL, and after 24 h of treatment cells were collected by gently scraping, and then pelleted by centrifugation at 400 *g* for 5 min at 4 °C. Proteomes were extracted, alkylated with IAA only, proteolyzed, and desalted as described above. Peptides were resolved by strong-cation exchange chromatography (SCX) using a Protein Pak Hi-Res SP 7 μm 4.6 × 100 mm column (Waters), using an Alliance liquid chromatograph (Waters) equipped with an automated fraction collector. Peptides were resuspended in 100% buffer A (5% ACN/0.1% FA) and loaded at 0.5 mL/min constant flow rate. Keeping column temperature at 30°C and flow rate at 0.5 mL/min, peptides were eluted using a segmented gradient of buffer B (1M potassium chloride (KCl) in 5% ACN/0.1% FA) as follows: 0–5 min (0% B), 5–35 min (0– 15% B), 35–50 min (15–50% B), 50–60 min (100% B), 60–75 min (0% B). In total 18 fractions were obtained, which were concentrated by vacuum centrifugation and desalted using solid-phase extraction with C18 MacroSpin columns (Nest Group). Peptides were lyophilized and stored at −80°C until MS analysis.

### Preparation of BSA/KEAP1 modified by itaconate

Bovine serum albumin (BSA) and recombinant human Kelch-like ECH-associated protein 1 (KEAP1) were dissolved in MS-grade H_2_O and mixed at equimolar amounts at a concentration of 2 μM. The mixture was reduced by 1 mM tris(2-carboxyethyl)phosphine) (TCEP) at room temperature for 1 h, and dialyzed against 100 mM ABC, 1 μM TCEP at 4°C overnight. The BSA/KEAP1 solution was divided into 3 equal fractions, and incubated at 4°C overnight with itaconate at 50, 5 and 0 mM concentrations, respectively. Itaconate-reacted proteins were concentrated by vacuum centrifugation, denatured using 6 M guanidinium chloride, 100 mM ABC, pH 8.4, and alkylated with IAA as described above. For LysC/trypsin digestion, the solution was diluted with 100 mM aqueous ABC, pH 8.4 to a final concentration of 0.6 M guanidinium chloride and digested using 1:50 w/w LysC at 37°C for 6 h. The solution was further diluted to 0.3 M guanidinium chloride, and digested using 1:25 w/w modified trypsin at 37°C overnight. For GluC or elastase digestion, the solution was diluted to 0.3 M guanidinium chloride, and digested using either 1:20 w/w GluC or elastase at 37°C overnight. Digested peptides were purified as described above.

### Liquid chromatography and peptide mass spectrometry

For analysis of proteomes reacted with iodoacetamide and acrylamide, the LC system consisted of a vented trap-elute configuration (Ekspert nanoLC 400, SCIEX) coupled to an Orbitrap Lumos mass spectrometer (Thermo Fisher Scientific, San Jose, CA) via a nano electro-spray DPV-565 PicoView ion source (New Objective). The trap column was fabricated with a 5 cm × 150 μm internal diameter silica capillary with a 2 mm silicate frit (Dhabaria et al. 2015), and pressure loaded with Poros R2-C18 10 μm particles (Life Technologies). The analytical column consisted of a 50 cm × 75 μm internal diameter column packed with ReproSil-Pur C18-AQ 3 μm particles (Dr. Maisch), and connected to a 3 cm electrospray emitter with 3 μm terminal inner diameter (Cifani et al. 2015). 5 μl of peptide solution were resolved over 150 min 3%–45% linear gradient of acetonitrile/ 0.1% formic acid (buffer B) in a water/0.1% formic acid (buffer A), delivered at 300 nL/minute. Precursor ions in the 400–2000 m/z range were filtered using the quadrupole and their spectra were recorded every 3 s using the Orbitrap (240,000 resolution), with an automatic gain control target set at 10^5^ ions and a maximum injection time of 100 ms. Data-dependent MS2 selection was enforced, limiting fragmentation to monoisotopic ions with charge 2–6, and dynamically excluding already fragmented precursors for 30 s (10 ppm tolerance). Selected precursors were isolated (Q1 isolation window 1.2 Th) and HCD fragmented (normalized stepped collision energy 24, 32, 40%) using the top speed algorithm. Fragmentation spectra were recorded in the Orbitrap at 50,000 resolution (AGC 50,000 ions, maximum injection time 86 ms).

For analysis of SCX-fractionated RAW264.7 proteomes, peptides were resolved using an Easy-nLC 1000 chromatograph (Thermo Fisher Scientific) in line with quadrupole-orbitrap MS (Q-Exactive HF, Thermo Fisher Scientific). Peptides were injected on a EasySpray reversed phase column (50 cm × 75 μm internal diameter) with integrated electrospray emitter (Thermo Fisher Scientific) and resolved with a constant flowrate of 300 nl/min using 2-45% linear gradient of acetonitrile in water (both containing 0.1% v/v formic acid) over 150 min, followed by a 45-90% gradient over 3 minutes and 25 minutes at constant 90% acetonitrile. Eluting peptides were transferred into the mass spectrometer via an EasySpray nano electro-spray ion source. Precursor ions in the 400-1500 m/z range were filtered using the quadrupole and recorded every 3 s using the Orbitrap (60,000 resolution, with 445.1200 ion used as lockmass), with an automatic gain control target set at 10^6^ ions and a maximum injection time of 50 ms. Data-dependent MS2 selection was enforced, limiting fragmentation to monoisotopic ions with charge 2– 5 and MS1 intensity greater than 5×10^4^, and dynamically excluding already fragmented precursors for 30 s (10 ppm tolerance). Selected precursors were isolated (Q1 isolation window 1.2 Th) and HCD fragmentated (normalized collision energy 30). Product ion spectra were recorded in the Orbitrap at 15,000 resolution (AGC 5×10^4^ ions, maximum injection time 54 ms).

BSA and KEAP1 reacted with itaconate peptides were resolved using an Easy-nLC 1000 chromatograph (Thermo Fisher Scientific) in line with quadrupole-Orbitrap MS (Fusion Lumos Orbitrap, Thermo Fisher Scientific). Peptides were loaded on a EasySpray reversed phase column (50 cm × 75 μm internal diameter) with integrated electrospray emitter (Thermo Fisher Scientific) and resolved with a constant flowrate of 300 nl/min using 2-45% linear gradient of acetonitrile in water (both containing 0.1% v/v formic acid) over 60 minutes, followed by a 45-90% gradient over 3 minutes and 25 minutes at constant 90% acetonitrile. Eluting peptides were transferred into the mass spectrometer via an EasySpray nano electro-spray ion source. Precursor ions in the 375–3000 m/z range were filtered using the quadrupole and recorded every 3 s using the Orbitrap (60,000 resolution, with 445.1200 ions used as lockmass), with an automatic gain control target set at 10^6^ ions and a maximum injection time of 50 ms. Data-dependent MS2 selection was enforced, limiting fragmentation to monoisotopic ions with charge 2– 5 and MS1 intensity greater than 5×10^4^, and dynamically excluding already fragmented precursors for 30 s (10 ppm tolerance). Selected precursors were isolated (Q1 isolation window 0.7 Th) and HCD fragmentated (normalized collision energy 30) using the top speed algorithm. Product ion spectra were recorded in the Orbitrap at 30,000 resolution (AGC 8×10^4^ ions, maximum injection time 54 ms).

### Data analysis

Raw files were converted to mgf open format using MSConvert within Proteowizard (Adusumilli & Mallick 2017) stable release 3.0. Spectra were matched to the reference human proteome as retrieved from UniProt (The UniProt Consortium 2017) on May 23rd 2019, supplemented with contaminant proteins from the cRAP database (Mellacheruvu et al. 2013) and decoy sequences generated using the PEAKS Studio suite v.8.0 (Bioinformatic Solution). Peptide spectral matching was performed using MSFragger v20190523 with FragPipe v9.3 and GUI v3.0 (Kong et al. 2017), or Byonic v2.7.84 (Bern et al. 2007; Bern & Kil 2011), or X!Tandem (version Alanine)(Craig & Beavis 2004) with SAMPEI. Precursor mass tolerance was set to 5 ppm for conventional searches (MSFragger “closed” search, Byonic without wildcard, and X!Tandem), and to 200 Da for agnostic PTM identification (MSFragger “open” search, Byonic with wildcard function enabled, and SAMPEI). All other search parameters were kept homogenous across searches, as permitted by the respective algorithms. Specifically, fragment ion mass tolerance was set to 20 ppm, trypsin with up to 3 missed cleavages was set as protease, M-oxidation and N/Q deamidation were set as variable modifications. All spectra were analyzed with Cys fixed modifications set to either carbamidomethylation or propionylation, dependent on the experimental conditions. PSMs were initially ranked by score (MSFragger hyperscore, Byonic score) and filtered to obtain a PSM level 1% FDR. Carbamidomethylated and propionylated cysteine-containing PSMs were then counted, considering both detected mass shifts corresponding to the specific adducts (i.e. +57 and +71 Da for carbamidomethylation and propionylation, respectively) and apparent mass shifts produced by the difference between variable and agnostically discovered PTM. In the case of carbamidomethylated Cys detected while setting propionylation as variable modification, a mass shift of −14 Da was used, corresponding to the difference between the observed 57 Da modification and the predicted 71 Da mass shift. In the case of propionylated Cys detected while setting carbamidomethylation as variable modification, a mass shift of +14 Da was used, corresponding to the difference between the observed 71 Da modification and the predicted 57 Da mass shift.

### Metabolomics

RAW264.7 cells were cultivated and treated as above. After 24 h of LPS treatment, metabolites were extracted and analyzed by liquid chromatography-MS (LC/MS) and gas chromatography-mass spectrometry (GC/MS, will edit once the data is available). For LC/MS, metabolites were extracted from cell pellets with ice-cold 80:20 methanol:water. After overnight incubation at −80**°**C, samples were vortexed and cleared by centrifugation at 20,000 *g* for 20 min. Supernatants were dried in a vacuum evaporator (Genevac EZ-2 Elite). Dried extracts were resuspended in in 40 μL of 97:3 water:methanol containing 10 mM tributylamine and 15 mM acetic acid. Samples were vortexed, incubated on ice for 20 min, and clarified by centrifugation at 20,000 G for 20 min at 4°C. LC-MS analysis used a Zorbax RRHD Extend-C18 column (150 mm × 2.1 mm, 1.8 μm particle size, Agilent Technologies). Solvent A was 10 mM tributylamine,15 mM acetic acid in 97:3 water:methanol, and solvent B was 10 mM tributylamine and 15 mM acetic acid in methanol, prepared according to the manufacturer’s instructions (MassHunter Metabolomics dMRM Database and Method, Agilent Technologies). LC separation was coupled to a 6470 triple quadrupole mass spectrometer (Agilent Technologies) which was operated in dynamic MRM scan type and negative ionization mode. Itaconate was identified at a retention time of ~13.4 minutes with an MRM transition of m/z 129 to 85.1 (primary transition used for quantitation), and m/z 129 to 41.3 (confirmatory). Retention time was also confirmed by injection of pure itaconate and the pure standard spiked into a pooled sample.

For GC/MS, metabolites were extracted from cell pellets with ice-cold 80:20 methanol:water containing 2 mM deuterated 2-hydroxyglutarate (D-2-hydroxyglutaric-2,3,3,4,4-d5 acid; deuterated-2HG) as an internal standard. After overnight incubation at −80°C, samples were vortexed and cleared by centrifugation at 21,000 G for 20 min at 4° C. Extracts were then dried in a vacuum evaporator (Genevac EZ-2 Elite). Dried extracts were resuspended by addition of 50 mL of methoxyamine hydrochloride (40 mg/mL in pyridine) and incubated at 30**°**C for 90 min with agitation. Metabolites were further derivatized by the addition of 80 mL of N-methyl-N-(trimethylsilyl) trifluoroacetamide (MSTFA) + 1% 2,2,2-trifluoro-N-methyl-N-(trimethylsilyl)-acetamide, chlorotrimethylsilane (TCMS; Thermo Scientific) and 70 mL of ethyl acetate (Sigma) and incubated at 37**°**C for 30 min. Samples were diluted 1:2 with 200 mL of ethyl acetate, then analyzed using an Agilent 7890A GC coupled to Agilent 5977 mass selective detector. The GC was operated in splitless mode with constant helium carrier gas flow of 1 mL/min and with a HP-5MS column (Agilent Technologies). The injection volume was 1 mL and the GC oven temperature was ramped from 60**°**C to 290**°**C over 25 min. Peaks representing compounds of interest were extracted and integrated using MassHunter vB.08.00 (Agilent Technologies) and then normalized to both the internal standard (deuterated-2HG) peak area and cell number or protein content as applicable. Ions used for quantification of metabolite levels were itaconate m/z 259 (confirmatory ion m/z 215) and deuterated-2HG m/z 252 (confirmatory ion m/z 354). Peaks were manually inspected and verified relative to known spectra for each metabolite. Pure itaconate (Sigma), both alone and spiked into a pooled sample, was used for confirmation.

### Western blot analysis

LPS-stimulated RAW264.7 cells were washed twice with PBS, resuspended in RIPA lysis buffer (150 mM NaCl, 50 mM Tris-HCl, 1% NP-40, 0.5% sodium deoxycholate, 0.1% SDS, pH 8.0), supplemented with protease inhibitor cocktail (0.5 mM AEBSF, 0.01 mM bestatin, 0.1 mM leupeptin, and 0.001 mM pepstatin). Cells were mechanically disrupted using S220 adaptive focused sonicator (Covaris) and the lysate was cleared by centrifugation at 15,000 *g* for 10 min. The protein concentration of the clarified lysates was determined using BCA assay according to the manufacturer’s instructions (Pierce). 30 μg of proteome were resolved by denaturing electrophoresis (SDS-PAGE) through a NuPAGE 10% Bis-Tris polyacrylamide gel (Invitrogen) with MOPS SDS running buffer, and electroeluted onto Immobilon FL polyvinylidene difluoride (PVDF) membrane (EMD Millipore). The membrane was blocked with 5% w/v non-fat dry milk for 1 h at room temperature and probed overnight at 4 °C with rabbit anti-IRG1 (1:1000, D6H2Y, Cell Signaling) and anti-β actin (1:20000, 13E5, Cell Signaling) antibodies. After washing with TBST buffer for 3 times, the membrane was then incubated with secondary antibody (donkey anti-rabbit, horseradish peroxidase (HRP) linked, 1:20,000) for 1 h at room temperature. Membranes were washed 5 times with TBST buffer and twice with TBS. Membranes were incubated 5 minutes at room temperature with SuperSignal west femto maximum sensitivity substrate (Pierce), and peroxidase activity was recorded using a BioRad imager.

## Supplementary Figures

**Supp. Fig. 1:**
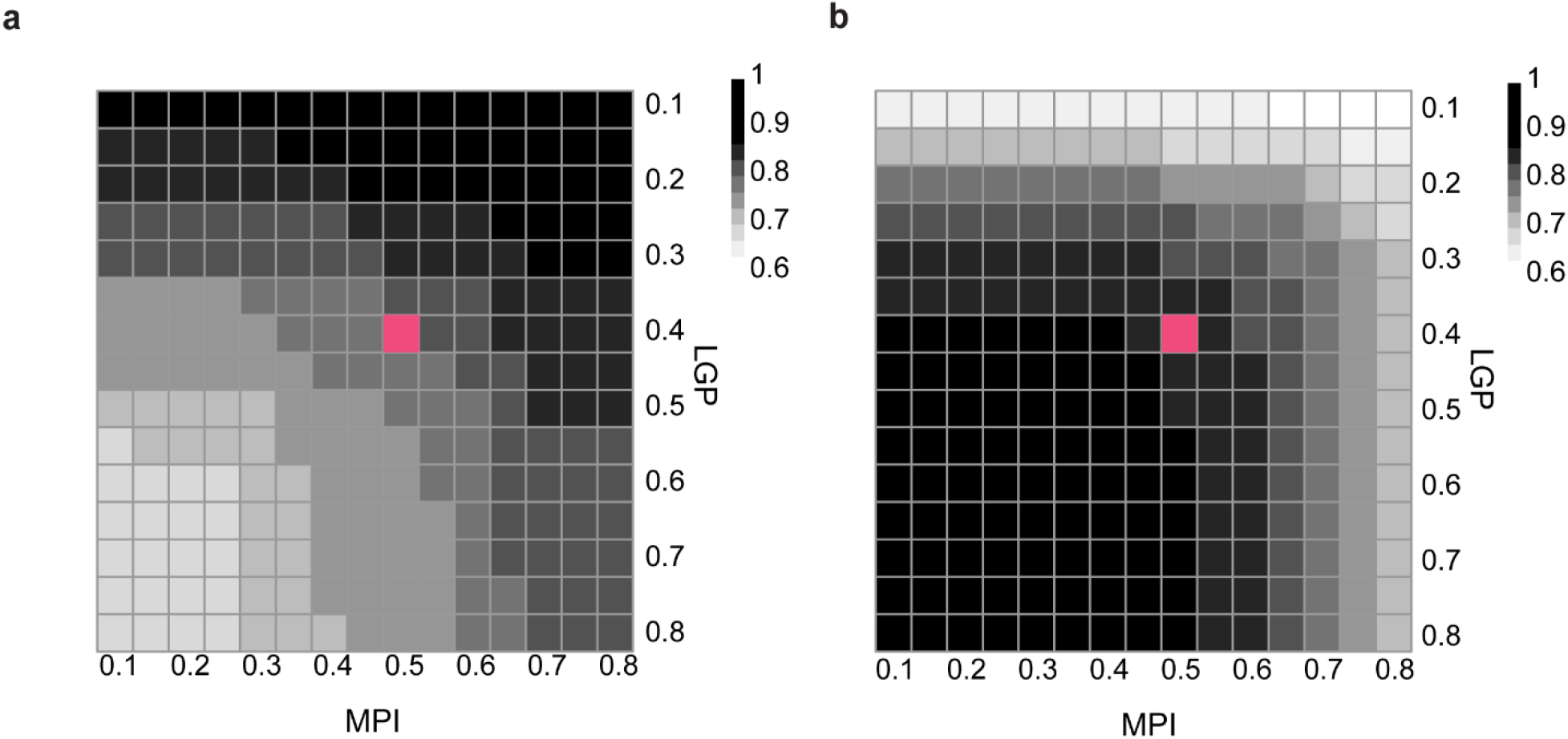
SAMPEI parameterization using specific differentially alkylated proteomes. Empirical optimization of MPI and LGP parameters to maximize precision (**a**) and recall (**b**), setting propionylation as a fixed modification. Red symbols denote the default user-specified values.

**Supp. Fig. 2:**
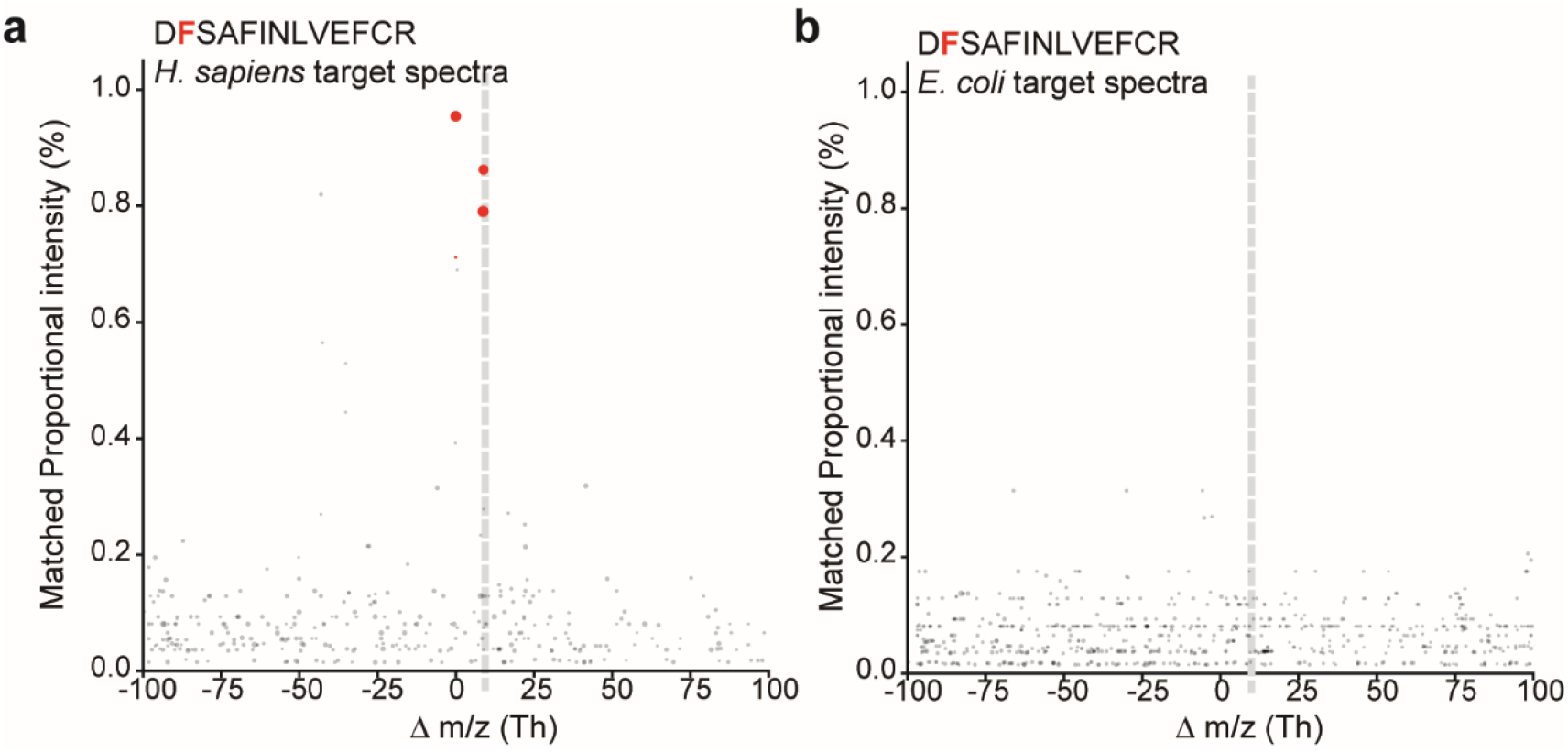
SAMPEI achieves sensitive and specific unbiased discovery of peptide modifications in complex cellular proteomes. a,b. Fraction of matched spectra as a function of modification mass shift (m/z) demonstrating accurate identification of synthetically fluorinated D(fluoro-F)SAFINLVEFCR peptide (17.99 Da mass shift, corresponding to 8.995 Th for +2 charged ions, as marked by dotted gray line) using as queries human cell extract containing the unmodified peptide (**a**) or an E. coli proteome not containing the unmodified chemoform, and thus serves as a negative control (**b**). Red symbols denote fluoro-peptide spectra matched by SAMPEI to the unmodified query PSM. Gray symbols denote matches of the fluorinated peptide with unrelated unmodified spectra.

**Supp. Fig. 3:**
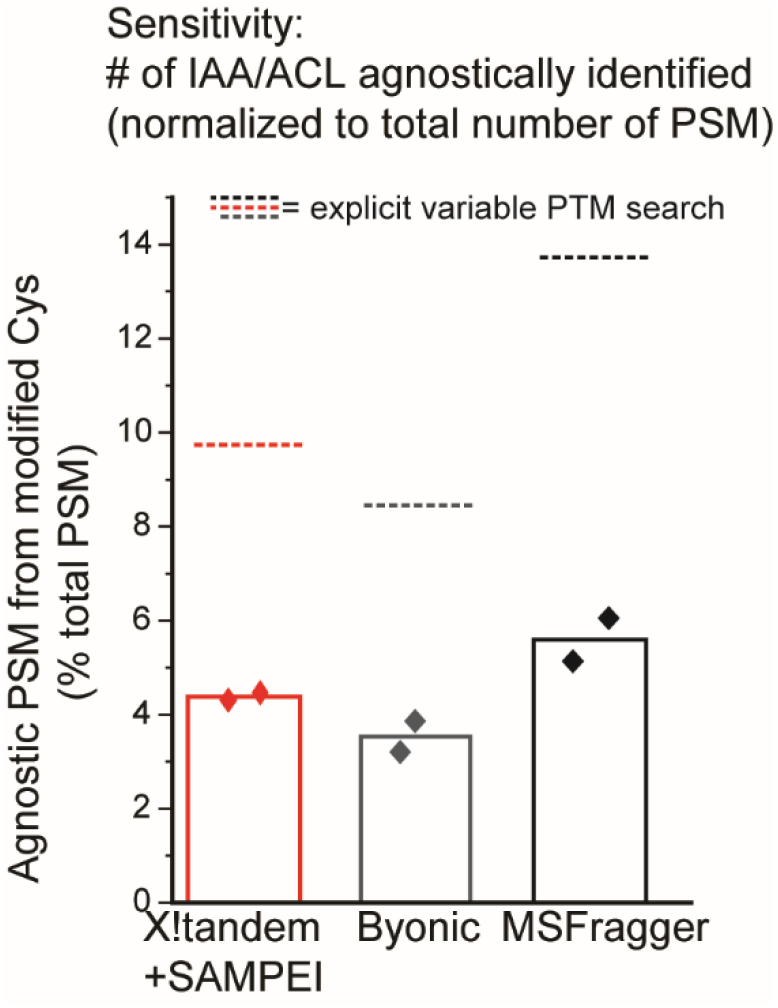
Sensitivity of SAMPEI, Byonic and MSFragger, as a fraction of total PSM. Recovery of cysteine alkylation by agnostic PTM analysis is measured as fraction of total PTM. Dotted lines indicate the mean fraction of modified cysteines detected with the relevant adducts as variable modification (Byonic, MSFragger), or by X!Tandem with SAMPEI.

**Supp. Fig. 4:**
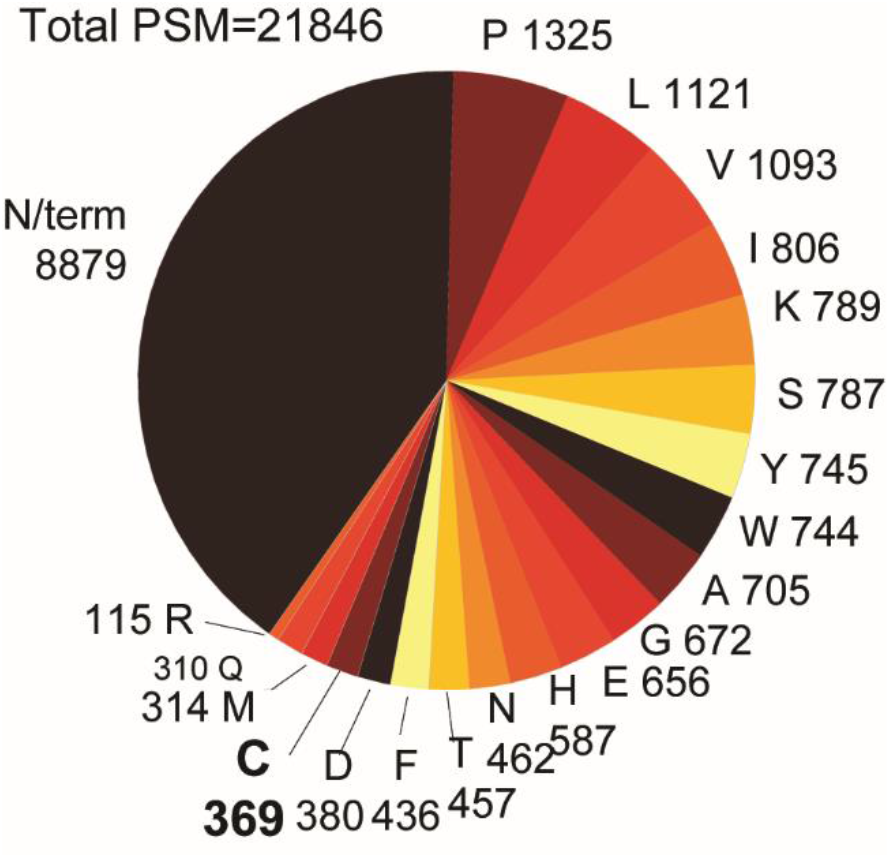
Distribution of putative modified sites for agnostically identified PTMs. Numerical values denote number of modified peptide-spectral matches.

**Supp. Fig. 5:**
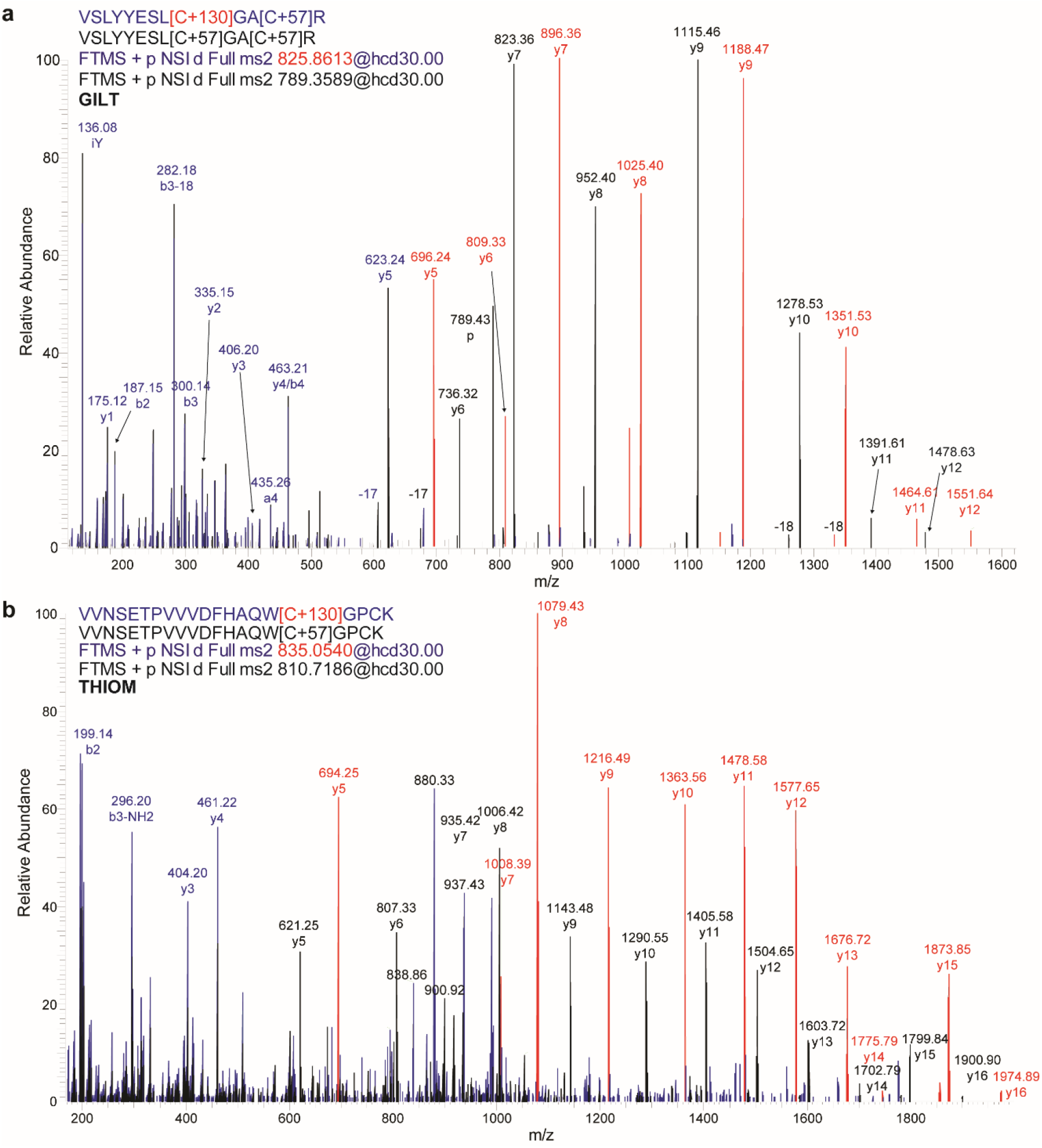
Representative mass spectra pairs for peptides with either unmodified experimentally induced +57 Da modification or biologic +130 Da modification of peptide cysteines. Representative mass spectra pairs of peptides with either unmodified cysteine (subject to experimental carbamidomethylation) or bearing +130 Da modifications detected in Gilt (**a**) and Thiom (**b**) proteins. High-resolution fragmentation mass spectra pairs were overlaid and annotated, showing diagnostic mass shift between modified peptides. Ions shared by both chemoforms are annotated in blue; ions solely matching the unmodified peptides are annotated in black; ions specific for the +130 Da modified peptides are annotated in red.

**Supp. Fig. 6:**
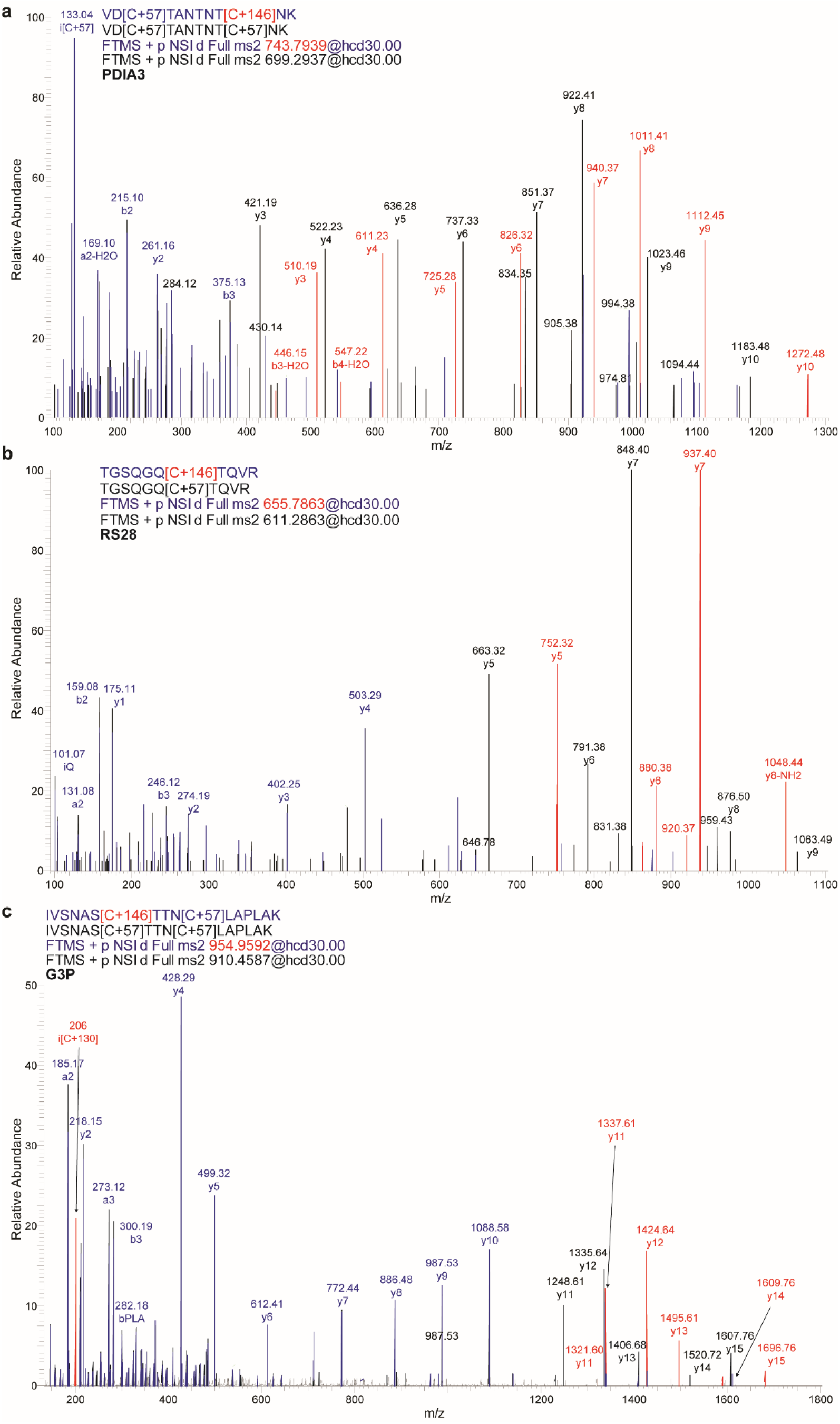
Representative mass spectra pairs for peptides with either unmodified experimentally induced +57 Da modification or biologic +146 Da modification of peptide cysteines. Representative mass spectra pairs of peptides with either unmodified cysteine (subject to experimental carbamidomethylation) or bearing +146 Da modifications detected in Pdia3 (**a**), Rs28 (**b**), and G3p (**c**) proteins. High-resolution fragmentation mass spectra pairs were overlaid and annotated, showing diagnostic mass shift between modified peptides. Ions shared by both chemoforms are annotated in blue; ions solely matching the unmodified peptides are annotated in black; ions specific for the +146 Da modified peptides are annotated in red.

**Supp. Fig. 7:**
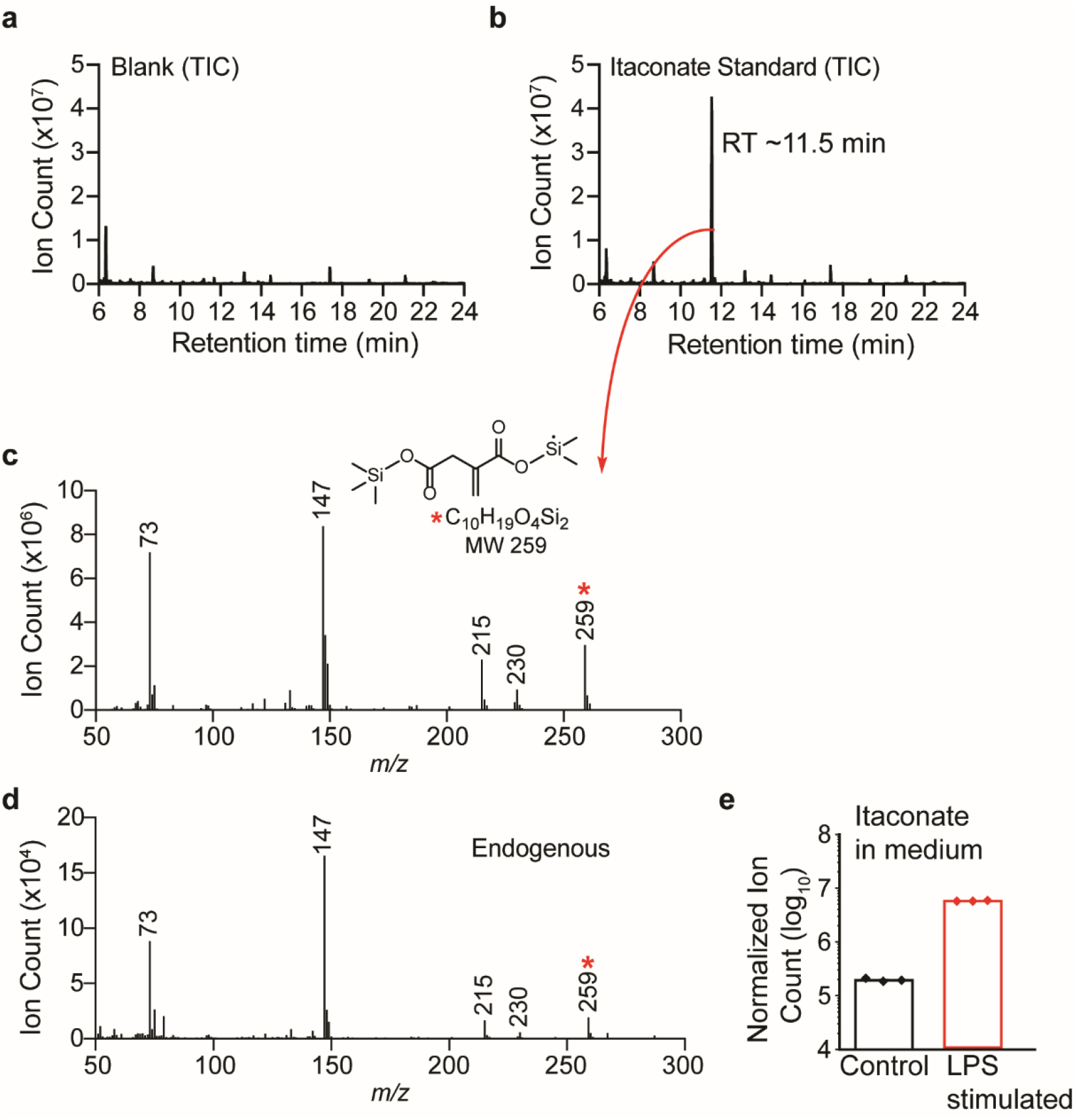
Quantification of itaconate production by LPS-stimulated macrophages. Total ion chromatograms (TIC) for blank solvent (**a**) or pure trimethylsilylated itaconate standard (**b**) measured by GC-MS (see Methods for details). The specific peak for trimethylsilyl-itaconate was detected at a retention time of ~11.5 minutes. **c.** Mass spectrum for the trimethylsilyl-itaconate peak at retention time 11.5 minutes; *m/z* 259 was used for subsequent itaconate quantification and *m/z* 215 as a confirmatory ion. The inset illustrates the structure of the 2,2,2-trifluoro-N-methyl-N-(trimethylsilyl)-acetamide chlorotrimethylsilane (TCMS)-derivatized itaconate ion with molecular weight (MW) 259. **d.** Quantification of itaconate production by RAW264.7 macrophages upon LPS stimulation as measured by LC-MS in medium (*n* = 3 biologic replicates).

**Supp. Fig. 8:**
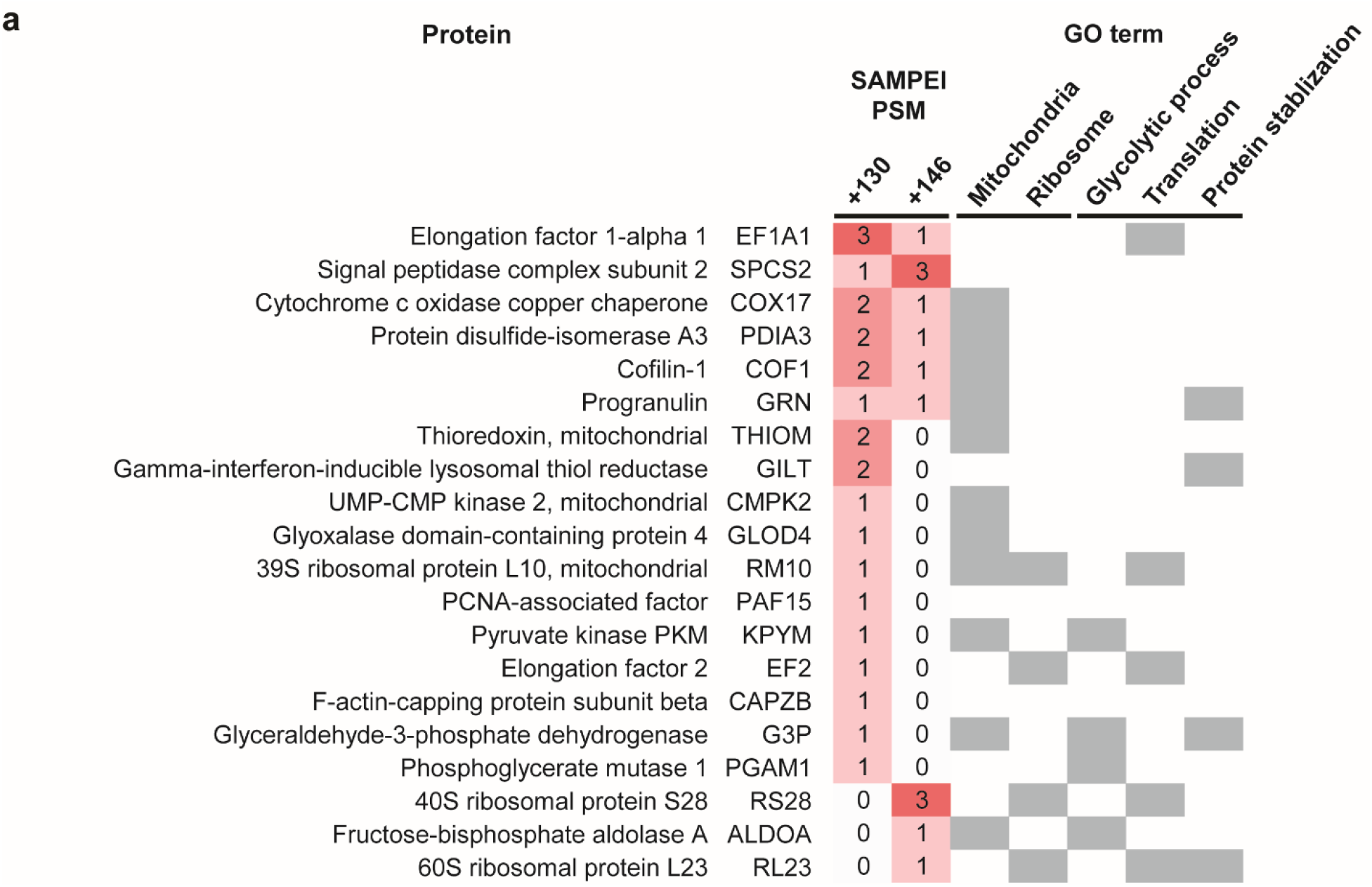
GO term enrichment of proteins with +130 Da and +146 Da peptide modifications.

**Supp. Fig. 9:**
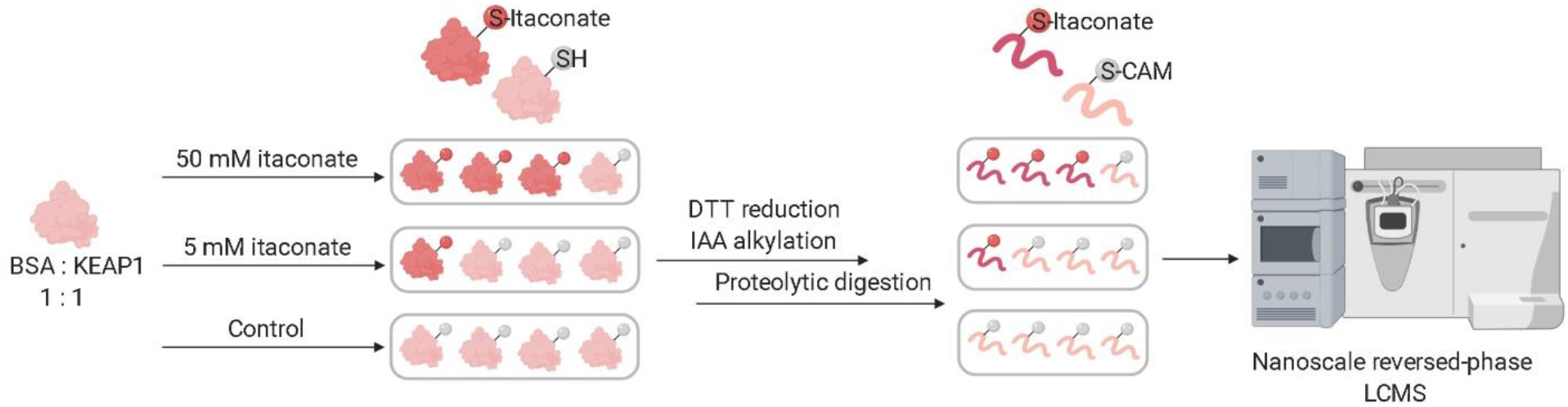
Schematic of itaconate reactions with purified BSA and KEAP1 proteins, followed by proteolysis and peptide chromatography and high-resolution tandem mass spectrometry.

**Supp. Fig. 10:**
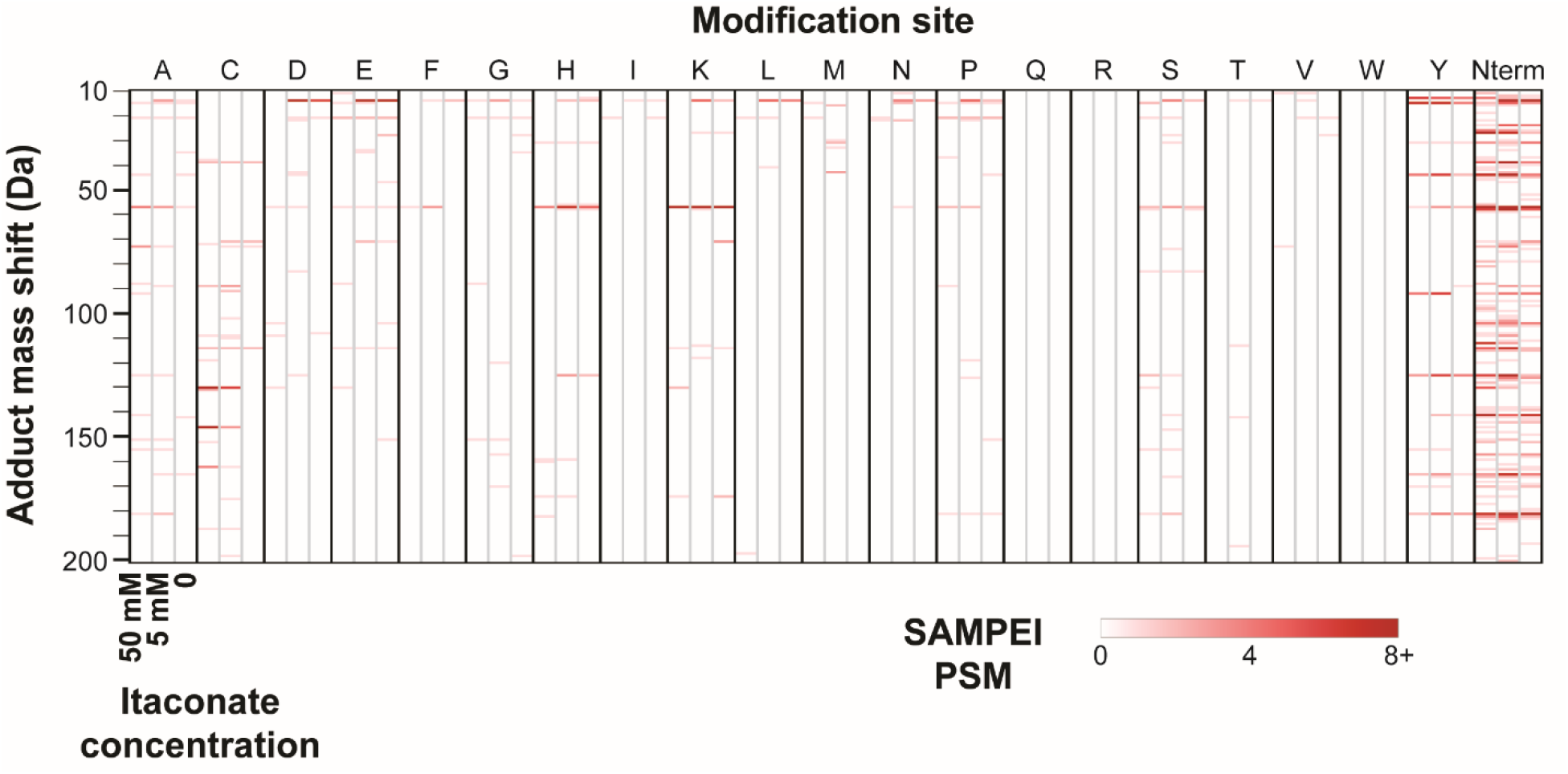
Agnostic PTM profiling of itaconate-reacted purified BSA and KEAP1 *in vitro*, as studied by tryptic peptide mass spectrometry. Frequency of PTM mass shifts of BSA and KEAP1 as a function of increasing concentrations of itaconic acid. For each putative localization, the three columns indicate adducts observed upon treatment with 50 mM (left column), 5 mM (middle) and 0 mM control itaconic acid (right column). Intensity of red marking indicates prevalence, expressed as number of SAMPEI PSMs.

**Supp. Fig. 11:**
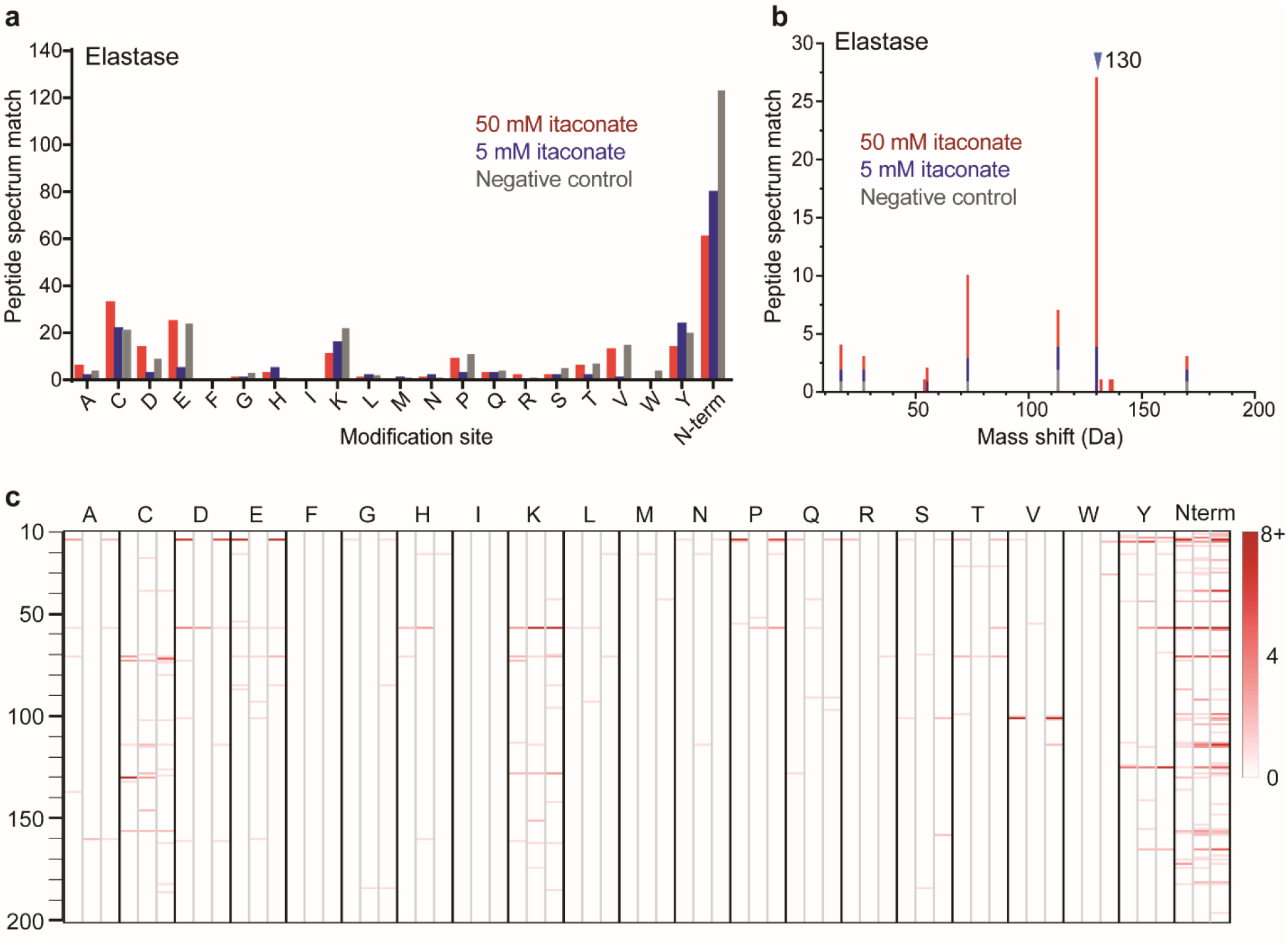
Agnostic PTM profiling of itaconate-reacted purified BSA and KEAP1 *in vitro*, as studied by elastase peptide mass spectrometry. **a.** Localization of non-canonical modifications of BSA and KEAP1, showing dependency of cysteine modifications on itaconate concentration. **b.** Frequency of observed adducts on BSA and KEAP1 treated with itaconate *in vitro* and negative control (as stacked column), showing dose-dependency of +130 Da adducts. **c.** Frequency of PTM mass shifts on BSA and KEAP1 amino acids treated with different doses of itaconic acid. For each localization, the three columns indicate adducts profiling obtained after treatment with 50 mM itaconate (left column), 5 mM itaconate (middle) and negative control (right column). Intensity of red marking indicates prevalence, expressed as number of SAMPEI PSMs.

**Supp. Fig. 12:**
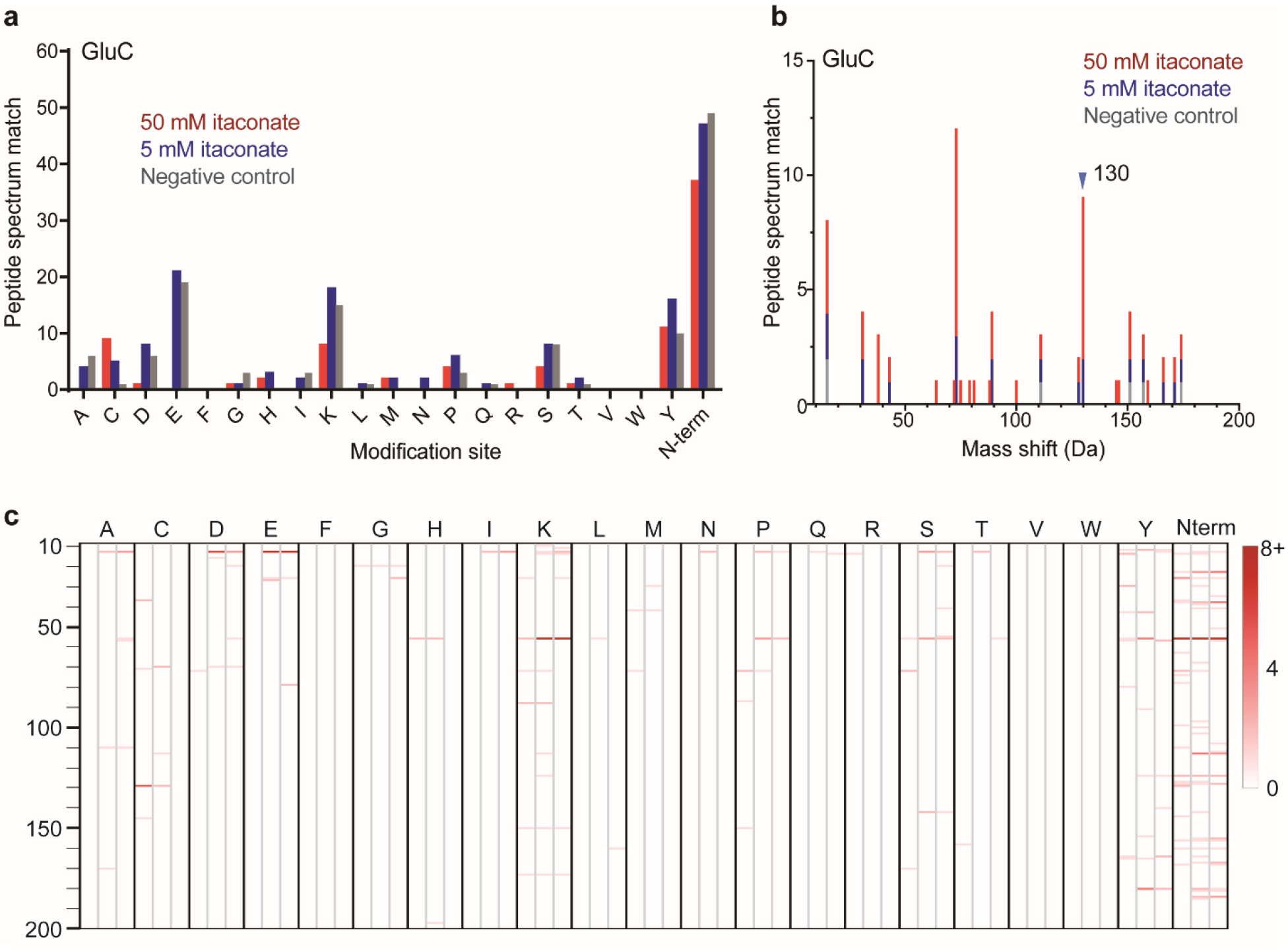
Agnostic PTM profiling of itaconate-reacted purified BSA and KEAP1 *in vitro*, as studied by GluC peptide mass spectrometry. **a.** Localization of non-canonical modifications BSA and KEAP1, showing dependency of cysteine modifications on itaconate concentration. **b.** Frequency of observed adducts on BSA and KEAP1 treated with itaconate *in vitro* and negative control (as stacked column), showing dose-dependency of +130 Da adducts. **c.** Frequency of PTM mass shifts on BSA and KEAP1 amino acids treated with different doses of itaconic acid. For each localization, the three columns indicate adducts profiling obtained after treatment with 50 mM itaconate (left column), 5 mM itaconate (middle) and negative control (right column). Intensity of red marking indicates prevalence, expressed as number of SAMPEI PSMs.

## Data availability

Mass spectrometry data have been deposited to the ProteomeXchange Consortium via the PRIDE (Perez-Riverol Y, et al. 2019) partner repository with the dataset identifiers PXD019793, PXD019827, PXD019858, and PXD019853. Processed files are available from Zenodo (http://doi.org/10.5281/zenodo.3899480).

## References

Adusumilli, R. & Mallick, P., 2017. Data Conversion with ProteoWizard msConvert. Methods in molecular biology (Clifton, N.J.), 1550(10), pp.339–368.

Aebersold, R. & Mann, M., 2003. Mass spectrometry-based proteomics. Nature, 422(6928), pp.198–207.

Aebersold, R. & Mann, M., 2016. Mass-spectrometric exploration of proteome structure and function. Nature, 537(7620), pp.347–355.

Aebersold, R. et al., 2018. How many human proteoforms are there? Nature chemical biology, 14(3), pp.206–214.

Alasoo, K. et al., 2015. Transcriptional profiling of macrophages derived from monocytes and iPS cells identifies a conserved response to LPS and novel alternative transcription. Scientific reports, 5(1), p.12524.

Bern, M., Cai, Y. & Goldberg, D., 2007. Lookup peaks: a hybrid of de novo sequencing and database search for protein identification by tandem mass spectrometry. Analytical Chemistry, 79(4), pp.1393–1400.

Bern, M.W. & Kil, Y.J., 2011. Two-dimensional target decoy strategy for shotgun proteomics. Journal of proteome research, 10(12), pp.5296–5301.

Bittremieux, W., Laukens, K. & Noble, W.S., 2019. Extremely Fast and Accurate Open Modification Spectral Library Searching of High-Resolution Mass Spectra Using Feature Hashing and Graphics Processing Units. Journal of proteome research, 18(10), pp.3792–3799.

Chi, H. et al., 2018. Comprehensive identification of peptides in tandem mass spectra using an efficient open search engine. Nature biotechnology, 36(11), p.1059.

Chick, J.M. et al., 2015. A mass-tolerant database search identifies a large proportion of unassigned spectra in shotgun proteomics as modified peptides. Nature biotechnology, 33(7), pp.743–749.

Cifani, P. et al., 2018. ProteomeGenerator: A Framework for Comprehensive Proteomics Based on de Novo Transcriptome Assembly and High-Accuracy Peptide Mass Spectral Matching. Journal of proteome research, 17(11), pp.3681–3692.

Cifani, P., Dhabaria, A. & Kentsis, A., 2015. Fabrication of Nanoelectrospray Emitters for LC-MS. Available at: http://www.nature.com/protocolexchange/protocols/3981.

Cordes, T. et al., 2016. Immunoresponsive Gene 1 and Itaconate Inhibit Succinate Dehydrogenase to Modulate Intracellular Succinate Levels. The Journal of biological chemistry, 291(27), pp.14274–14284.

Cox, J. & Mann, M., 2008. MaxQuant enables high peptide identification rates, individualized p.p.b.-range mass accuracies and proteome-wide protein quantification. Nature biotechnology, 26(12), pp.1367–1372.

Craig, R. & Beavis, R.C., 2004. TANDEM: matching proteins with tandem mass spectra. Bioinformatics (Oxford, England), 20(9), pp.1466–1467.

Creasy, D.M. & Cottrell, J.S., 2002. Error tolerant searching of uninterpreted tandem mass spectrometry data. Proteomics, 2(10), pp.1426–1434.

Creasy, D.M. & Cottrell, J.S., 2004. Unimod: Protein modifications for mass spectrometry. Proteomics, 4(6), pp.1534–1536.

Dasari, S. et al., 2010. TagRecon: high-throughput mutation identification through sequence tagging. Journal of proteome research, 9(4), pp.1716–1726.

Devabhaktuni, A. et al., 2019. TagGraph reveals vast protein modification landscapes from large tandem mass spectrometry datasets. Nature biotechnology, 37(4), pp.469–479.

Dhabaria, A., Cifani, P. & Kentsis, A., 2015. Fabrication of Capillary Columns with Integrated Frits for Mass Spectrometry. Available at: http://www.nature.com/protocolexchange/protocols/3925.

Eng, J.K., McCormack, A.L. & Yates, J.R., 1994. An approach to correlate tandem mass spectral data of peptides with amino acid sequences in a protein database. Journal of the American Society for Mass Spectrometry, 5(11), pp.976–989.

Everett, L.J., Bierl, C. & Master, S.R., 2010. Unbiased statistical analysis for multi-stage proteomic search strategies. Journal of proteome research, 9(2), pp.700–707.

Guijas, C. et al., 2018. METLIN: A Technology Platform for Identifying Knowns and Unknowns. Analytical Chemistry, 90(5), pp.3156–3164.

Gut, P. & Verdin, E., 2013. The nexus of chromatin regulation and intermediary metabolism. Nature, 502(7472), pp.489–498.

Han, X. et al., 2011. PeaksPTM: Mass spectrometry-based identification of peptides with unspecified modifications. Journal of proteome research, 10(7), pp.2930–2936.

Hokao, R. & Francia, G., 2001. Differential display. Methods in molecular medicine, 57, pp.297–305.

Huang, X. et al., 2013. ISPTM: an iterative search algorithm for systematic identification of post-translational modifications from complex proteome mixtures. Journal of proteome research, 12(9), pp.3831–3842.

Jensen, O.N., Podtelejnikov, A.V. & Mann, M., 1997. Identification of the components of simple protein mixtures by high-accuracy peptide mass mapping and database searching. Analytical Chemistry, 69(23), pp.4741–4750.

Jeong, J. et al., 2011. Novel oxidative modifications in redox-active cysteine residues. Molecular & cellular proteomics : MCP, 10(3), p.M110.000513.

Kahnert, K. et al., FLEXIQuant-LF: Robust Regression to Quantify Protein Modification Extent in Label-Free Proteomics Data. bioRxiv, DOI: 10.1101/2020.05.11.088492

Kamal, A.H.M. et al., 2018. Inflammatory Proteomic Network Analysis of Statin-treated and Lipopolysaccharide-activated Macrophages. Scientific reports, 8(1), p.164.

Keller, A. et al., 2002. Empirical statistical model to estimate the accuracy of peptide identifications made by MS/MS and database search. Analytical Chemistry, 74(20), pp.5383–5392.

Kentsis, A., 2005. Between light and eye: Goethe’s science of color and the polar phenomenology of nature. ArXiv:physics/0511130

Kong, A.T. et al., 2017. MSFragger: ultrafast and comprehensive peptide identification in mass spectrometry-based proteomics. Nature methods, 14(5), pp.513–520.

Kulkarni, R.A. et al., 2019. A chemoproteomic portrait of the oncometabolite fumarate. Nature chemical biology, p.1.

Lepack, A.E. et al., 2020. Dopaminylation of histone H3 in ventral tegmental area regulates cocaine seeking. Science, 368(6487), pp.197–201.

MacCoss, M.J. et al., 2002. Shotgun identification of protein modifications from protein complexes and lens tissue. Proceedings of the National Academy of Sciences of the United States of America, 99(12), pp.7900–7905.

Mann, M. & Jensen, O.N., 2003. Proteomic analysis of post-translational modifications. Nature biotechnology, 21(3), pp.255–261.

Mann, M. & Wilm, M., 1994. Error-tolerant identification of peptides in sequence databases by peptide sequence tags. Analytical Chemistry, 66(24), pp.4390–4399.

Mellacheruvu, D. et al., 2013. The CRAPome: a contaminant repository for affinity purification-mass spectrometry data. Nature methods, 10(8), pp.730–736.

Michalski, A., Cox, J. & Mann, M., 2011. More than 100,000 detectable peptide species elute in single shotgun proteomics runs but the majority is inaccessible to data-dependent LC-MS/MS. Journal of proteome research, 10(4), pp.1785–1793.

Mills, E.L. et al., 2018. Itaconate is an anti-inflammatory metabolite that activates Nrf2 via alkylation of KEAP1. Nature, 556(7699), pp.113–117.

Na, S. & Paek, E., 2009. Prediction of novel modifications by unrestrictive search of tandem mass spectra. Journal of proteome research, 8(10), pp.4418–4427.

Nielsen, M.L., Savitski, M.M. & Zubarev, R.A., 2006. Extent of modifications in human proteome samples and their effect on dynamic range of analysis in shotgun proteomics. Molecular & cellular proteomics : MCP, 5(12), pp.2384–2391.

Paulsen, C.E. & Carroll, K.S., 2013. Cysteine-mediated redox signaling: chemistry, biology, and tools for discovery. Chemical reviews, 113(7), pp.4633–4679.

Perez-Riverol Y, et al., 2019. The PRIDE database and related tools and resources in 2019: improving support for quantification data. Nucleic Acids Res 47(D1):D442–D450.

Poole, L.B. & Nelson, K.J., 2008. Discovering mechanisms of signaling-mediated cysteine oxidation. Current Opinion in Chemical Biology, 12(1), pp.18–24.

Qin, W. et al., 2019. S-glycosylation-based cysteine profiling reveals regulation of glycolysis by itaconate. Nature chemical biology, 15(10), pp.983–991.

Qin, W., Yang, F. & Wang, C., 2020. Chemoproteomic profiling of protein-metabolite interactions. Current Opinion in Chemical Biology, 54, pp.28–36.

Rattigan, K.M. et al., 2018. Metabolomic profiling of macrophages determines the discrete metabolomic signature and metabolomic interactome triggered by polarising immune stimuli. A. Ahmad, ed. PloS one, 13(3), p.e0194126.

Savitski, M.M., Nielsen, M.L. & Zubarev, R.A., 2006. ModifiComb, a new proteomic tool for mapping substoichiometric post-translational modifications, finding novel types of modifications, and fingerprinting complex protein mixtures. Molecular & cellular proteomics : MCP, 5(5), pp.935–948.

Searle, B.C. et al., 2004. High-throughput identification of proteins and unanticipated sequence modifications using a mass-based alignment algorithm for MS/MS de novo sequencing results. Analytical Chemistry, 76(8), pp.2220–2230.

Seim, G.L. et al., 2019. Two-stage metabolic remodelling in macrophages in response to lipopolysaccharide and interferon-γ stimulation. Nature metabolism, 1(7), pp.731–742.

Seo, Y.H. & Carroll, K.S., 2009. Profiling protein thiol oxidation in tumor cells using sulfenic acid-specific antibodies. Proceedings of the National Academy of Sciences of the United States of America, 106(38), pp.16163–16168.

Sihvola, V. & Levonen, A.-L., 2017. Keap1 as the redox sensor of the antioxidant response. Archives of biochemistry and biophysics, 617, pp.94–100.

Smith, L.M., Kelleher, N.L. Consortium for Top Down Proteomics, 2013. Proteoform: a single term describing protein complexity. Nature methods, 10(3), pp.186–187.

Steen, H. & Mann, M., 2004. The ABC“s (and XYZ”s) of peptide sequencing. Nature Reviews Molecular Cell Biology, 5(9), pp.699–711.

Strelko, C.L. et al., 2011. Itaconic acid is a mammalian metabolite induced during macrophage activation. Journal of the American Chemical Society, 133(41), pp.16386–16389.

The UniProt Consortium, 2017. UniProt: the universal protein knowledgebase. Nucleic acids research, 45(D1), pp.D158–D169.

West, A.P. et al., 2011. TLR signalling augments macrophage bactericidal activity through mitochondrial ROS. Nature, 472(7344), pp.476–480.

Wilhelm, M. et al., 2014. Mass-spectrometry-based draft of the human proteome. Nature, 509(7502), pp.582–587.

Zhang, D. et al., 2019. Metabolic regulation of gene expression by histone lactylation. Nature, 574(7779), pp.575–580.

Zheng, Q. et al., 2020. Non-enzymatic covalent modifications: a new link between metabolism and epigenetics. Protein & cell, 9(39), p.23.

Zhou, F. et al., 2013. Genome-scale proteome quantification by DEEP SEQ mass spectrometry. Nature communications, 4, p.2171.

Zolg, D.P. et al., 2018. ProteomeTools: Systematic Characterization of 21 Post-translational Protein Modifications by Liquid Chromatography Tandem Mass Spectrometry (LC-MS/MS) Using Synthetic Peptides. Molecular & cellular proteomics : MCP, 17(9), pp.1850–1863.

